# The functional evolution of collembolan Ubx on the regulation of abdominal appendage formation

**DOI:** 10.1101/2023.12.01.564554

**Authors:** Yan Liang, Yun-Xia Luan

## Abstract

*Folsomia candida* is a tiny soil-living arthropod belonging to the Collembola, which is an outgroup to Insecta. It resembles insects as having a pair of antennae and three pairs of thorax legs, while it also possesses three abdominal appendages: a ventral tube located in the first abdominal segment (A1), retinaculum in A3 and furca in A4. Collembolan Ubx and AbdA specify abdominal appendages, but they are unable to repress appendage marker gene *Dll*. The genetic basis of collembolan appendage formation and the mechanisms by which Ubx and AbdA regulate *Dll* transcription and appendage development remain unknown. In this study, we analysed the developmental transcriptomes of *F. candida* and identified candidate appendage formation genes, including *Ubx* (*FcUbx*). Expression data revealed the dominance of *Dll* over *Ubx* during the embryonic 3.5 and 4.5 days, suggesting Ubx is deficient in suppressing *Dll* at early appendage formation stages. Furthermore, via electrophoretic mobility shift assays and dual luciferase assays, we found that the binding and repression capacity of FcUbx on *Drosophila Dll* resembles those of the longest isoform of *Drosophila* Ubx (DmUbx_Ib), while the regulatory mechanism of the C-terminus of FcUbx on *Dll* repression is similar to that of crustacean *Artemia franciscana* Ubx (AfUbx), demonstrating that the function of collembolan Ubx is intermediate between that of Insecta and Crustacea. In summary, our study provides novel insights into collembolan appendage formation and sheds light on the functional evolution of Ubx. Additionally, we propose a model that collembolan Ubx regulates abdominal segments in a context-specific manner.

## Introduction

Extant arthropods are traditionally classified into four major groups: chelicerates, myriapods, crustaceans, and hexapods (including proturans, collembolans, diplurans, and insects) (Budd & Telford, 2009; Giribet et al., 2001; Hughes & Kaufman, 2002b). Throughout their evolution, arthropods gradually reduce abdominal appendages, particularly insects, which entirely lack these appendages in their adult stages (Jockusch & Smith, 2015; Matsuda, 2017). The establishment of a body plan is regulated by a cascade of orchestrated transcription factors during embryogenesis, including maternal effect genes, pair-rule genes, gap genes and Hox genes (Peel et al., 2005). Hox genes encode transcription factors characterized by a helix-turn-helix DNA-binding homeodomain and play pivotal roles in the identification and appendage formation of each segment along the body axis of arthropods (Angelini & Kaufman, 2005a, 2005b; Budd & Telford, 2009; Hughes & Kaufman, 2002a). Typically, Hox genes are organized in a cluster on the chromosome and exhibit both spatial and temporal collinearity in their expression patterns (Durston et al., 2011). *Dll* (*Distal-less*) serves as a marker gene for the appendage primordium in a wide variety of animals (Panganiban et al., 1997), within some groups of pancrustaceans (including crustaceans and hexapods) (Browne & Patel, 2000; Cohen et al., 1991; Jockusch et al., 2004), it acts as a downstream target of Ubx (*Ultrabithorax*) (Buffry et al., 2023; Cohen, 1990; Galant & Carroll, 2002; Gebelein et al., 2002; Palopoli & Patel, 1998; Ronshaugen et al., 2002). In *Drosophila melanogaster*, the Ubx gene generates six isoforms (*Ib*, *Ia*, *IIb*, *IIa*, *IVb*, *IVa*) via alternative splicing events, with variation in the architecture of the linker region between the YPWM motif and homeodomain (HD domain) (Geyer et al., 2015; Passner et al., 1999; Reed et al., 2010). Specifically, the *Ib* isoform (DmUbx_Ib) has the longest linker region, and the *IVa* isoform (DmUbx_IVa) does not contain the linker. Functional assays demonstrated that the longest linker is indispensable for the suppression of *Dll* expression (Gebelein et al., 2002); in addition, the QAQA domains and poly-Ala stretch located at the C-terminus are also crucial for the repression function (Galant & Carroll, 2002; Gebelein et al., 2002; Hughes & Kaufman, 2002b; Ronshaugen et al., 2002). In general, these structures facilitate Ubx the ability to bind to the enhancer of *Dll* (Dll304) and consequently suppress the expression of the *Dll* gene (Cohen et al., 1993; Gebelein et al., 2002). In contrast, in branchiopod crustacean *Artemia franciscana*, Ubx (AfUbx) loses its capacity to repress *Dll* expression (Ronshaugen et al., 2002). This functional shift can be attributed to the presence of phosphorylation sites within the C-terminus of AfUbx, which impede its ability to suppress limb development (Galant & Carroll, 2002; Ronshaugen et al., 2002). To date, the evolutionary processes and the molecular mechanism that give rise to the loss of abdominal appendages in adult hexapods remain elusive.

Collembola (springtails) is a group of basal hexapods, whose phylogenetic position is intermediated between aquatic crustaceans and terrestrial insects (Gao et al., 2008; Luan et al., 2005; Timmermans et al., 2008). Collembolans bear three distinct types of abdominal appendages (M. T. Fountain & S. P. Hopkin, 2005): the ventral tube (or collophore) in abdominal segment 1 (A1), which is involved in osmoregulation with the external environment; the retinaculum in A3, a structure for holding the springing organ furca in place; and the furca in A4, a robust jumping apparatus (Tully & Potapov, 2015). A previous study demonstrated that the collembolan Ubx specifies A1 and AbdA determines A4, and they both regulate the morphological formation of A2 and A3 (Konopova & Akam, 2014). Intriguingly, collembolan Ubx, AbdA and *Dll* are concurrently expressed in the collembolan abdomen during the embryonic stage, indicating that collembolan Ubx and AbdA does not repress *Dll* expression in abdominal segments (Palopoli & Patel, 1998). However, the molecular mechanisms of how collembolan Ubx and AbdA interact with *Dll* and thus regulate appendage formation are unclear.

To address this question, we conducted a computational analysis of the developmental transcriptomes of Collembola, *F. candida* (Denmark strain), covering embryos and juveniles to adults. Through this analysis, we identified the genes potentially involved in appendage formation during embryogenesis, with *Ubx* being a notable inclusion. Expression data indicated that *Ubx* could not suppress *Dll* in the early appendage formation stages. Furthermore, we investigated the transcriptional regulatory functions of Ubx. Our findings revealed that the binding capacity and repression activity of *F. candida* Ubx (FcUbx) on *D. melanogaster Dll* are analogous to those in *D. melanogaster* Ubx (DmUbx_Ib) (Galant & Carroll, 2002; Gebelein et al., 2002; Ronshaugen et al., 2002); and the regulatory mode and site(s) in the C-terminus of FcUbx resemble those of Ubx in the branchiopod crustacean *A. franciscana* (AfUbx) (Galant & Carroll, 2002; Ronshaugen et al., 2002). Based on these data, we depicted a molecular functional evolutionary model of Ubx from crustaceans to basal hexapods to insects in the broad scope of panarthropods and proposed that the collembolan Ubx might exert its repression function in distinct abdominal segments in a context-specific manner.

## Materials and methods

### Sample collection and transcriptome sequencing

First, *F. candida* (Denmark strain) was synchronized for three generations: some adult individuals (F0 generation) were transferred to a new Petri dish (10 cm), a container with a layer of a mixture of plaster of Paris and activated charcoal at a ratio of 9:1 by weight (Krogh, 2009). After 24 hours of oviposition, the eggs (F1 generation) were subsequently transferred to a new Petri dish for one month of culture to reach sexual maturity. The procedure was repeated twice to obtain synchronized adults of the F3 generation. Approximately 300 adults of the F3 generation were then transferred to three new Petri dishes for 24 hours of oviposition, and all eggs were transferred to a new culture dish. Next, all eggs laid by the F3 generation were collected and transferred to new culture dishes (5 cm) every 24 hours, except on the last day, when eggs were collected at 12 hours after the adults had been transferred.

After 51 days of collection and culture, we obtained 51 samples ranging from day 0.5 to day 50.5. Fifteen developmental samples, including eggs (from days 0.5 to 9.5), juveniles (days 12.5, 19.5 and 28.5), and adults (days 31.5 and 45.5), were chosen for transcriptome sequencing. RNA was prepared and extracted separately from 15 samples by using a miRNeasy mini kit (QIAGEN, Germany) according to the manufacturer’s instructions. The transcriptomes were sequenced on the BGISEQ-500 platform by BGI (Yuzuki, 2015) (Shenzhen, China). All of the raw reads had been published (BioProject PRJNA433725) (Liang et al., 2019).

### Transcriptome analysis and gene annotation

The reads mapping and differential gene expression analysis were identical to the previous study (Liang et al., 2019). The genome we used was non-chromosome-level genomic data from the published *F. candida* genome (Luan et al., 2023). The genome index was built using bowtie2 (Langmead & Salzberg, 2012), and the reads were mapped using tophat2 (Kim et al., 2013) The transcript abundance (RPKM) was estimated using cufflinks (Trapnell et al., 2012), with the gene annotation file from the whole genome. Gene Ontology (GO) annotations were accomplished by running BLAST2GO (Conesa et al., 2005) against the UniProt database (UniProt release 2018_02) (Bairoch et al., 2005). The expression profile was obtained from the data matrix representing the expression abundance (RPKM, reads per kilobase of transcript per million fragments mapped) from 15 transcriptomes. The distribution of gene expression was visualized as Ridgeline Plots in ‘ggplot2’ (Wickham, 2016). Additionally, to explore the relationships among developmental stages, we employed several analyses. Principal component analysis was conducted using the “prcomp” function, while Pearson correlation analysis was executed using the “cor” function in R. For hierarchical clustering analysis, we first scaled the dataset using the “Euclidean” method and then applied the “hclust” function with the “ward.D2” method. Data visualization was achieved using “ggplot2” and “pheatmap” (Kolde, 2012) in RStudio (2023.3.0.386) (RStudio, 2020).

### Mining of genes involved in appendage formation

To identify genes associated with the appendages formation from time-series bulk RNA-seq data, we employed the Short Time-series Expression Miner (STEM) approach (Ernst & Bar-Joseph, 2006; Ernst et al., 2005), which attempts to assign genes to previously defined temporal trajectory/development (Bar-Joseph et al., 2012). Through the embryonic observation, PCA analysis, and hierarchical clustering, we selected four key time points: day 1.5 (E_1.5d, blastula, no appendages), day 3.5 (E_3.5d, early stage of appendage formation), day 5.5 (E_5.5d, mid-stage of appendage formation) and day 7.5 (E_7.5d, mature stage of appendage formation) for mining genes were involved in appendage formation. STEM software was used to classify all the clusters according to the abundance of gene expression. Subsequently, genes in the clusters that were supposed to correlate with appendage formation were sorted out and then annotated by Blast2GO software (Conesa et al., 2005).

### Expression profiling of Hox genes and *Dll* in *D. melanogaster* and *F. candida*

The gene expression profile of *D. melanogaster* was downloaded from flybase (http://ftp.flybase.org/releases/current/precomputed_files/genes/gene_rpkm_matrix_fb _2023_05.tsv.gz) (Brown et al., 2014). The expression values of Hox genes and *Dll* were subset from the expression profile of *D. melanogaster* and *F. candida*. Notably, the *Ubx* in *F. candida* was fragmented into five transcripts (XLOC_011518, XLOC_011661, XLOC_011777, XLOC_011778 and XLOC_011779) (Supplementary Data 1). To calculate the expression of *Ubx*, we computed and accumulated the RPKM of those transcripts. The normalization of gene expression was applied in two approaches. Normalized RPKM, the original RPKM was normalized by comparing the minimum and maximum of each gene throughout the embryonic stages to elucidate the expression pattern and the lowest and highest expression stage of a gene. Z-score, the original RPKM was normalized by the z-scale in R (RStudio, 2020), which can reflect the expression pattern of all the selected genes.

### Mining of genes correlated with *Ubx* from transcriptomes

To predict the putative function or the underlying regulation of Ubx, based on the gene expression profile, we used the Spearman correlation algorithm (Spearman correlation coefficient r > 0.9, *p* < 0.01) to mine genes coexpressed with *Ubx* during embryogenesis. The coexpressed genes were extracted and then annotated by Blast2GO of gene ontology terms (GO terms) and KEGG pathway annotation (Conesa et al., 2005; Kanehisa & Goto, 2000).

### Acquisition of *Ubx*, *Exd* and *Hth* sequences

The complete sequences of two isoforms of *Ubx* in *F*. *candida* (Collembola) and the partial sequences from *Sinentomon erythranum* (Protura) and *Campodea augens* (Diplura) were cloned by degenerate PCR from the homeodomain, further obtained by 5’ RACE and 3’ RACE. Specifically, two isoforms of FcUbx were validated by using exon-intron and exon-exon junction PCR with overlapping primers spanning the linker region. *Exd* and *Hth* were identified via BLAST (Altschul et al., 1990; McGinnis & Madden, 2004) from transcriptomes and further validated by PCR cloning. All sequences were submitted to NCBI (OR593736, OR593737, OR593738, OR593739, OR604006, OR604007). The sequences of Panarthropod Ubx were aligned by MAFFT (Rozewicki et al., 2019), and the gaps were removed manually. The multiple sequence alignment was visualized in Jalview (Waterhouse et al., 2009).

### Protein expression and purification

The N-terminus truncated collembolan proteins, FcU1, FcU2, Exd and Hth used for protein expression were constructed in the pGEX-4t-1 plasmid (GST-tag inserted) and transformed into *E*. *coli* competent cells (BL21 strain). Protein expression was first started from 500 mL of Luria–Bertani (LB) medium (pH = 7) cultured at 37 °C at 220 rpm. Until it reached the exponential growth phase (OD600 = 0.8∼1.0), isopropyl β-D- 1-thiogalactopyranoside (IPTG) was added (0.1 mM) to 2 L of LB medium. The whole culture was then subjected to low-temperature induction of protein expression at 16 °C and 220 rpm for 12 hours. All the GST-tagged proteins were purified in native conditions according to the manufacturer’s instructions (Sangon Biotech, Shanghai, China). All proteins were quantitated by comparing them to a set of bovine serum albumin (BSA) concentration gradients: 750, 500, 250, 125, 100, 75, 50, 25, and 0 ng/μl by Coomassie blue staining and further confirmed by anti-GST western blotting.

### Electrophoretic mobility shift assays (EMSAs)

We identified a putative FcDll Element (PFE) in *F. candida* through a search within the approximately 4000 bp intergenic genomic region upstream of the first exon of the collembolan *Dll* gene. This search utilized the Hox binding consensus sequence (5’- TATA-3’) (Berger et al., 2008; Ekker et al., 1991; Noyes et al., 2008) and the Exd binding consensus sequence (5’-TGAT-3’) (Ryoo et al., 1999; Slattery et al., 2011).

For EMSAs, the DMXR element (Gebelein et al., 2002; Gebelein et al., 2004), a repression regulatory element of *Dll* (Dll304) in *D. melanogaster*, and the putative regulatory element of *Dll* in *F. candida* (Putative FcDll Element, PFE) were utilized as a DNA probe in each assay separately. The probe was first synthesized as two separate primers (forward and reverse strands), which were tagged with Cy5 at the 5’ end. The primers were diluted to a concentration of 1 μM. Subsequently, 50 μl of each primer was mixed and incubated at 95 °C and then cooled to room temperature. The primers were annealed, resulting in the formation of double-stranded DNA probes. 20 ng of DNA probe was used in each EMSA, and the amount of protein used in each EMSA was 10 pmol of GST, 0.2, 1, and 1.5 pmol of FcU1, 0.4, 2, and 3 pmol of FcU2 and 1 pmol of Exd and Hth. The procedures for electrophoretic mobility shift assays (EMSAs) were adjusted and visualized according to the protocol (He et al., 2016). The 20 μl EMSA solution: 5× EMSA buffer: Tris (0.1 M), glycerol (25%), and BSA (0.2 mg/ml).

2.5 M MgCl2, 1 M DTT, double-distilled water, poly(dI-dC), protein, and Cy5-labelled DNA probe. The reaction was incubated for 25 minutes at 25 °C. The 12% non-denaturing polyacrylamide gel contained double-distilled water, 5× TBE buffer, 50% glycerol, polyacrylamide (the monomer: dimer ratio was 80:1), 10% ammonium persulfate, and tetramethyl ethylenediamine (TEMED). Electrophoresis was carried out on ice, with the blank gel pre-electrophoresed at 120 V for 1 hour. Subsequently, 20 μl of the reaction mixture was loaded into each vial of the gel, and the samples were run at 150 V for 1-1.5 hours. For reaction visualization, the entire gel with the glass container was scanned directly with the A Starion FLA-900 phosphorimager (Fujifilm, Japan).

### *Drosophila* S2 cell transfection and dual luciferase reporter assays

The complete sequences of *D. melanogaster* Ubx (DmUbx_Ib, DmUbx_IVa) and collembolan Ubx (FcU1, FcU2), the truncated collembolan Ubx (FcU1/°C, FcU2/°C) and chimeric Ubx of *Drosophila* and collembolan (Dm/Fc_L, Dm/Fc_C) were cloned into the pAC5.1 expression vector. To test the repression on DMXR, this regulatory sequence was constructed into a pGL3-promoter vector (Promega), which contains the SV40 enhancer and firefly luciferase. Transfection experiments were carried out using the Qiagen Effectene reagent (Qiagen, Germany), according to the manufacturer’s protocol. Initially, *Drosophila* S2 cells were cultured in Schneider’s *Drosophila* medium (Sigma‒Aldrich, USA) supplemented with 10% fetal bovine serum (HyClone, USA). After 24 hours, the cells were aliquoted into 48-well plates at 150 μl per well. After 12 hours, 50 μl of DNA-enhancer mixture, containing 1.2 μl of enhancer (the reagent from the Effectene reagent kit), 0.075 μg of pGL3-promoter vector, 0.075 μg of pAC5.1 vector, and 0.2 μl of the Renilla fluorescent vector (pRL) were added to the mixture, followed by an 8-minute incubation. Then, 4 μl of Effectene and 200 μl of cell culture medium were added, and the transfected cells were cultured for 48 h in a 27 °C incubator. Each protein vector was set with technical triplicates.

The Dual luciferase reporter assay kit (Promega, USA) was used to measure the reporter gene expression, and the luciferase activities were detected by a Modulus^TM^ Microplate Luminometer (Turner BioSystems, USA). The repression activity of each protein is represented by the average ratio of Firefly: Renilla luciferase activity. To estimate the relative repression of the proteins on the DMXR, we compared the repression activity of each sample with that of DmUbx_Ib. For pairwise comparisons, we performed Student’s t-test in R (RStudio, 2020).

## Results

### Development and time-course transcriptomes of *F. candida*

The embryos of *F. candida* (Denmark strain) took approximately 10 days to reach the juvenile stage when incubated at 21 °C (Michelle T. Fountain & Steve P. Hopkin, 2005). Referring to the observation of the embryonic developmental stages in *F. candida* (Shanghai strain) (Gao et al., 2006), the samples selected for sequencing are illustrated in Figure 1A. The embryos in the 0-0.5 day (E_0.5d) predominantly ranged from the 4- cell stage to the blastula stage. By 1.5 days, most embryos had progressed in the gastrula stage. In 2.5 days, the initial phase of tissue differentiation stage, characterized by the segmentation and formation of appendage primordia. At 3.5 days, the antenna and thorax appendages started segmentation and elongation; the furca was observed by 4 days (Gao et al., 2006). During the period of 4.5 to 6.5 days, the middle phase of the tissue development stage progressed, accompanied by the growth and maturation of appendages. From 7.5 to 8.5 days, in the late phase of tissue differentiation, the appendages were fully formed, and the animals were preparing for hatching. Finally, at 9.5 days, the animals were actively moving within the eggshell, and some individuals had already hatched. Typically, it took these juveniles one month to reach sexual maturity as adults (Figure 1A, 1C).

**Figure 1.**
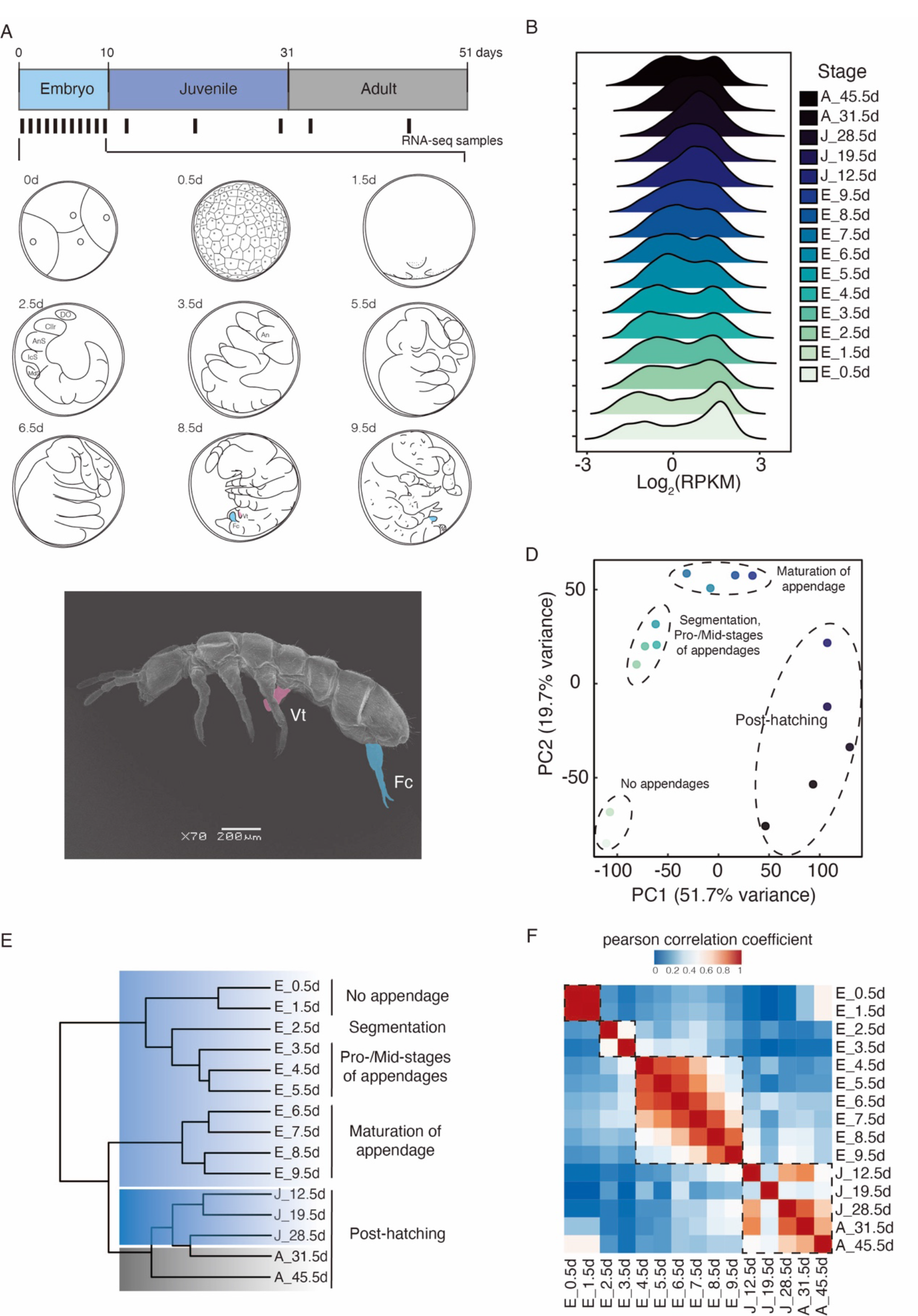
Analysis of developmental transcriptomes of *F*. *candida*. **A**. Schematic of samples for developmental transcriptome sequencing and the observation of embryonic development. Whole-animal collections were made from embryos (blue), juveniles (grey blue) and adults (grey). The bars indicate that different developmental samples were collected. 0d (d, days after oviposition): 4-cell stage; 0.5d: blastula stage; 1.5d: gastrula stage; 2.5d: Initial phase of tissue differentiation stage; 3.5-6.5d: middle phase of tissue differentiation stage; 7.5-8.5d: Late phase of tissue differentiation stage; 9.5d: prehatching stage. An: antenna; AnS: antenna segment; Cllr: clypeolabrum; DO: dorsal organ; Fc: furca (in blue colour); IcS: intercalary segment; MdS: mandibular segment; Vt: ventral tube (in pink colour). **B**. Distribution of gene expression of 25,038 genes from 15 developmental transcriptomes. RPKM, reads per kilobase per million, normalizes the raw count by transcript length and sequencing depth. **C**. Scanning electron microscopy (SEM) of the whole animal of adult *F. candida*. Vt: ventral tube (shade in pink colour), Fc: furca (shade in blue colour). **D**. Principal component analysis of the developmental transcriptomes. The dashed ellipses indicate the stages of appendage development. **E**. Simplified hierarchical clustering of all developmental transcriptomes. The stages of appendage development are shown on the right. **F**. Heatmap of the Pearson correlation coefficient of all the samples; the dashed rectangles indicate the relationship of samples.

The analysis of developmental transcriptomes recaptured the expression of a total of 25,803 genes throughout the developmental process (Figure 1B, Supplementary Data 1). Transcriptomic analyses elucidated that the relationships of all the developmental samples were consistent with the observation of development (Figure 1A, 1D, 1E, 1F). In general, all the post-hatching samples (juveniles and adults) clustered together, demonstrating that the hatching event acted as a critical developmental transitional event (Figure 1D, 1E, 1F). The embryonic samples were grouped separately: in the early stages E_0.5d and E_1.5d, no appendages were observed, and these two stages were grouped; the embryonic stages E_2.5d and E_3.5d, particularly marked by the segmentation and formation of appendage primordia, were clustered; and the period spanning from E_4.5d to E_9.5d, deemed the development of appendages, formed a distinct group. Specifically, E_4.5d to E_5.5d represent the mid-phase of appendage formation, and E_6.5d to E_9.5d were identified as the maturation of appendage formation (Figure 1D, 1E, 1F).

### Mining of genes related to appendage development in Collembola

Springtails bear three pairs of thoracic legs and appendages in the abdomen (Figure 1C). These abdominal appendages, however, exhibit distinct morphological characteristics from each other (Figure 1C). To identify genes associated with the establishment of appendages, especially the abdominal appendages, we conducted a Short Time-series Expression Miner analysis (STEM analysis) (Ernst & Bar-Joseph, 2006), which would assign genes to one of several previously defined developmental trajectories (Bar-Joseph et al., 2012). Based on the embryonic observation (Figure 1A) and transcriptomic analyses (Figure 1D, 1E, 1F), we deem the formation of appendages as four stages for our analysis: E_1.5d, corresponding to the blastula stage without any appendage structure; E_3.5d, representing the early stage of appendage development, marked by the emergence of the appendage primordia; E_5.5d, indicating middle stage of appendage development; and E_7.5d, the late stage of appendage development.

The number of expressed genes (RPKM > 0) in these four stages was 15,131, 18,693, 19,114 and 20,841 genes, respectively (Figure 2A, Supplementary Data 1, 2). The STEM analysis automatically categorized these genes into 50 profiles, among which 12 clusters were significantly statistically enriched (permutation test, *p* < 0.001). Per the morphological observations (Figure 1A) and the developmental trend we defined, the emergence of appendages is characterized by gradual development, with the transition of gene expression levels from low to high (Figure 2B). Cluster 42, which reflected this trend, was selected for further investigation (Figure 2C). Cluster 42 contained a total of 865 genes and 2,318 transcripts, of which 1,229 were annotated by Blast2GO (Supplementary Data 3). These genes were annotated as 10 biological processes of GO terms (Figure 2D) and involved in 32 KEGG pathways (Supplementary Data 3). In particular, 66 transcripts were associated with the developmental process, of which 36 transcripts were annotated, and 18 genes are the morphogenesis and appendages-related genes (Table 1, Supplementary Data 3). These genes constitute a set of conventional hierarchical developmental genes (Peel et al., 2005), including the body patterning gene *Notch* (de Celis et al., 1998; Kurata et al., 2000; O’Day, 2006; Rauskolb & Irvine, 1999) and Engrailed (Brower, 1986; van de Heuvel et al., 1993); limb proximal-distal axis formation genes *Exd, Hth* (Wu & Cohen, 1999); and the body plan “selector genes”, the Hox genes *Ubx*, *Antp*, and *Scr* (Weatherbee & Carroll, 1999). This cluster also comprised genes relevant to muscle and actin function, *e.g.*, *MYO1E* (*Myo61F*, orthologous in *D. melanogaster*) is an unconventional myosin-like protein (McIntosh & Ostap, 2016), *Lhx1* (*Lim1,* orthologous in *D. melanogaster*), which regulates tarsal segments (Natori et al., 2012); and *FBN2* (*CG7526*, orthologous in *D. melanogaster*) involves in fibrosis (Mead et al., 2022).

**Figure 2.**
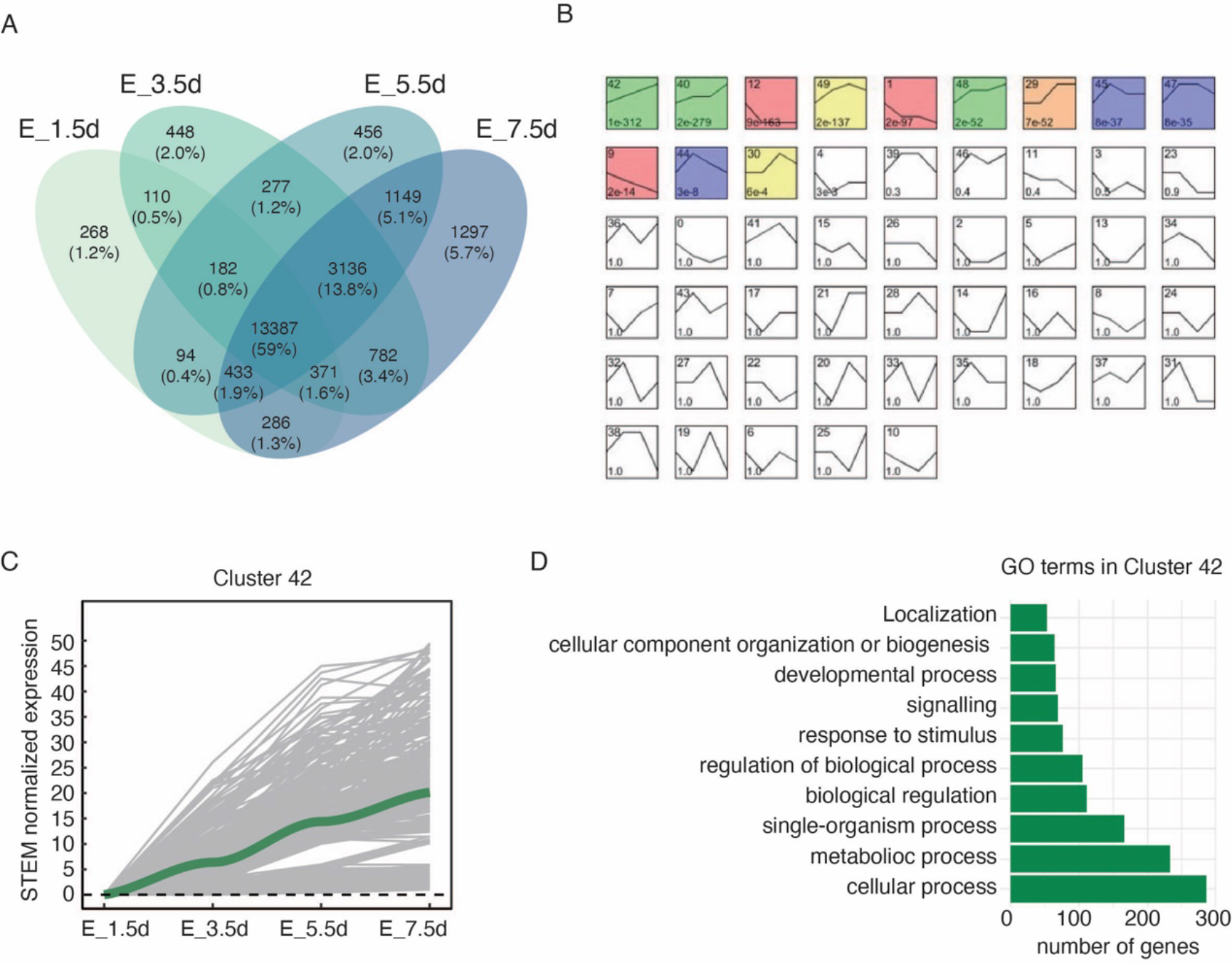
Identification of genes involved in appendage formation. **A.** Venn diagram illustrating the number and percentage of expressed genes of each intersection among the four appendage formation stages. **B.** A screenshot of different gene expression profiles identified by short time-series expression miner (STEM) analysis. The number of each profile is shown at the upper left in the square. The coloured profiles indicated that genes were clustered significantly (permutation test, *p* < 0.001). **C.** The STEM normalized expression of Cluster 42. Values higher than 50 were excluded. **D.** Annotated GO terms of Cluster 42.

**Table 1.** Genes classified in morphogenesis by BLAST2GO from Cluster 42.

| Gene_ID | Gene_symbol | Orthologous in<br>*D. melanogaster* | Molecular function |
| --- | --- | --- | --- |
| XLOC_013891 | FBN2 | CG7526 | embryonic limb<br>morphogenesis |
| XLOC_001350 | Exd | Exd | somatic muscle development |
| XLOC_002645 | Actin | Actin | muscle contraction |
| XLOC_017923 | BMI1 | Psc/Su(z)2 | segment specification |
| XLOC_019142 | GATA-3 | pnr/grn | anatomical structure<br>morphogenesis |
| XLOC_016846 | Wnt16 | Wnt5/ Wnt4 | Wnt signaling pathway |
| XLOC_022735 | En | engrailed | trunk segmentation |
| XLOC_011518 | Ubx | Ubx | specification of segmental<br>identity |
| XLOC_011519 | Antp | Antp | specification of segmental<br>identity |
| XLOC_015540 | ROR1 | Ror | inner ear development |
| XLOC_005027 | FOX | FoxF | anatomical structure<br>morphogenesis |
| XLOC_013406 | MYO1E | Myo61F/ Myo31DF/<br>CG15831 | microtubule-based<br>movement |
| XLOC_016945 | Notch | Notch | anterior/posterior pattern<br>specification |
| XLOC_011524 | Scr | Scr | specification of segmental<br>identity |
| XLOC_009469 | Ski | CG7233/Snoo | embryonic limb<br>morphogenesis |
| XLOC_019633 | Hth | Hth | segmentation |
| XLOC_007689 | LARGE2 | No Data | muscle cell cellular<br>homeostasis |
| XLOC_016637 | Lhx1 | Lim1 | anatomical structure<br>morphogenesis |

In summary, STEM analysis effectively identified several genes closely associated with appendage formation, thus validating the predefined stages utilized for identifying appendage formation genes.

### *Dominance* of *Dll* over *Ubx* during appendage formation in *F. candida*

In general, the Hox cluster is characterized by spatial (Figure 3A) and temporal collinearity (Monteiro & Ferrier, 2006). However, previously reported genome assemblies of *F. candida* (Faddeeva-Vakhrusheva et al., 2017; Luan et al., 2023) reveal that *Scr*, *Antp* and *Ubx* inserted into the anterior region of the Hox complex, indicating a lack of spatial collinearity (Figure 3A) (Faddeeva-Vakhrusheva et al., 2017). Our expression analysis of Hox genes in *D. melanogaster* also reveals a lack of temporal collinearity (Figure 3B), consistent with the previous study (Gaunt, 2015). Similarly, the Hox genes in *F. candida* do not display temporal collinearity (Figure 3D). For instance, the *pb* gene was transcribed at E_3.5d, occurring later than *Dfd* and *Scr*, and the most posterior gene *AbdB* was highly expressed at the early stage of E_4.5d; Notably, *Ubx* specifically displayed the highest expression levels at the appendage maturation stage E_7.5d.

**Figure 3.**
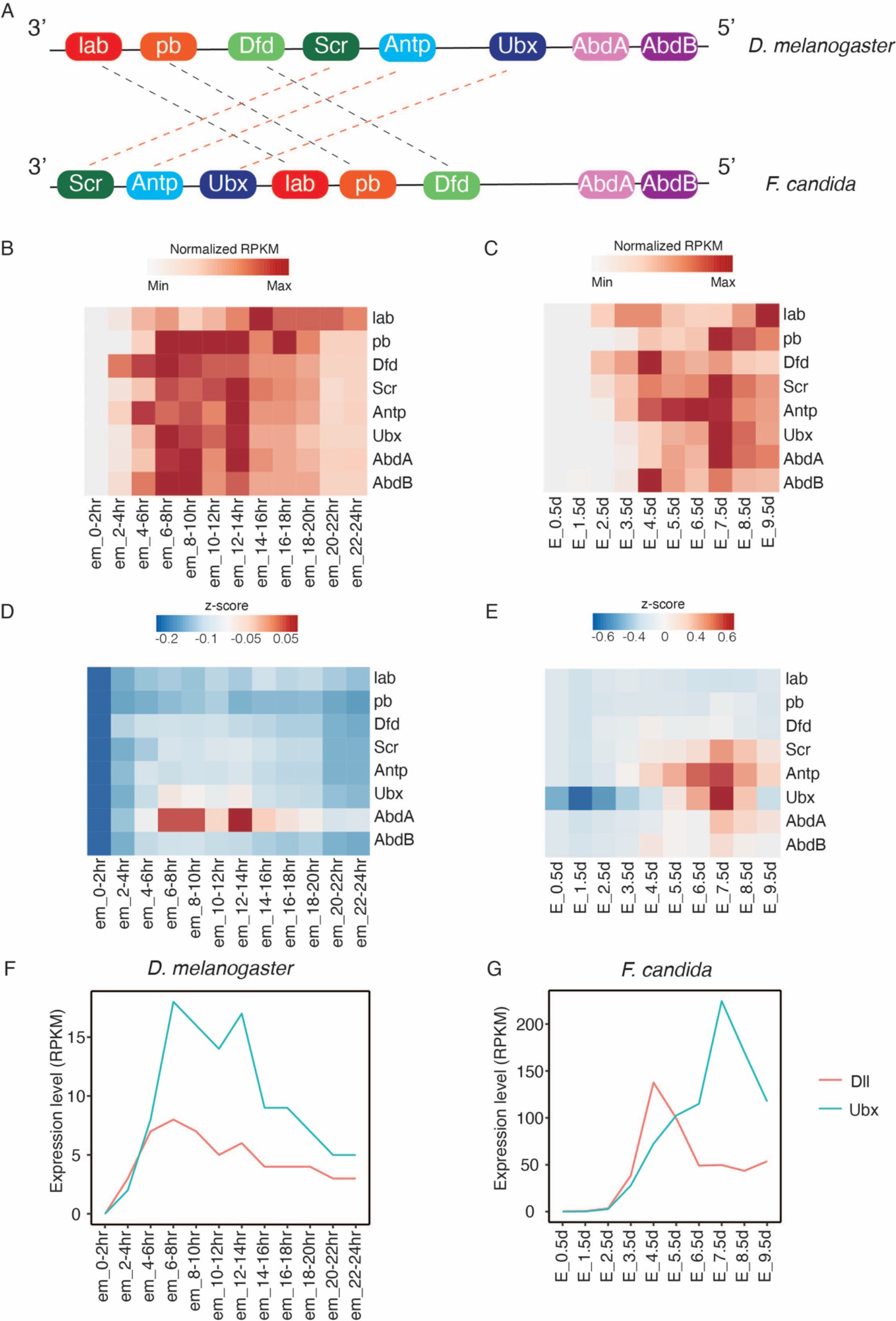
Comparison of the Hox cluster and *Dll* gene expression between *D. melanogaster* and *F. candida*. **A.** Genomic architecture of the Hox cluster in *D. melanogaster* (Refined from Pearson, et al. 2005) and *F. candida* (Refined from Faddeeva-Vakhrusheva, et al. 2017). The distances between genes were not scaled in proportion to the original genomic distance. Dashed lines indicate that the Hox genes are rearranged. **B, C, D, E, F, G.** Expression profiles of Hox genes and *Dll* in *D. melanogaster* **(B, D, F)** and *F. candida* **(C, E, G)**. Normalized RPKM, the RPKM were normalized by the minimum and maximum expression value of each gene throughout embryonic stages. Z-score, the RPKM values were normalized by z-scale.

In *D. melanogaster*, Ubx acts as a *Dll* repressor during embryogenesis in the abdominal segments (Gebelein et al., 2002). The transcriptomic expression data demonstraes that Drosophila *Ubx* exhibited higher expression levels than *Dll* (Figure 3F) during embryogenesis, suggesting its role in repressing *Dll*. Conversely, in *F. candida*, *Dll* predominated over *Ubx* during the E_3.5d and E_4.5d stages (Figure 3G), indicating that FcUbx may not repress *Dll* during the appendage formation stage. However, from the E_5.5d stage onward, *Ubx* expression surpassed that of *Dll*, indicating its potential to regulate or suppress *Dll* during the appendage maturation.

### Collembolan Ubx can bind to Hox/Exd/Hth DNA binding motifs

Ubx regulates *Dll* through DNA binding and transcriptional repression (Galant & Carroll, 2002; Gebelein et al., 2002; Grenier & Carroll, 2000; Ronshaugen et al., 2002). However, the mechanism of collembolan Ubx interacting with *Dll* remains unclear. To address this, we first obtained two complete isoforms of *Ubx* from *F. candida*, which are produced by alternative splicing and vary at the linker region (FcUbx, isoform 1, FcU1 with a linker of GQSYL; isoform 2, FcU2 without a linker) (Supplementary Data 4); and investigated their binding capacity and repression activity through *in vitro* and *in vivo* assays.

We conducted the electrical mobile shift assays (EMSAs) to examine the binding capacity of collembolan Ubx on an exogenous *Dll* element, the *Dll* regulatory element of *D. melanogaster* (DMXR, the repression element on Dll304) (Gebelein et al., 2004) (Figure 4A). The proteins of two isoforms of collembolan Ubx (FcUbx1, FcUbx2) can readily bind to DMXR, demonstrating that FcUbxs exhibit binding capability and that their linker region does not affect this binding ability (Figure 4B). Furthermore, the dimer Exd/Hth stimulates Ubx binding to DNA and forms Ubx/Exd/Hth trimeric protein complexes (Figure 4B), thereby demonstrating the ability of collembolan Ubx/Exd/Hth complexes to bind exogenous *Dll* in vitro.

**Figure 4.**
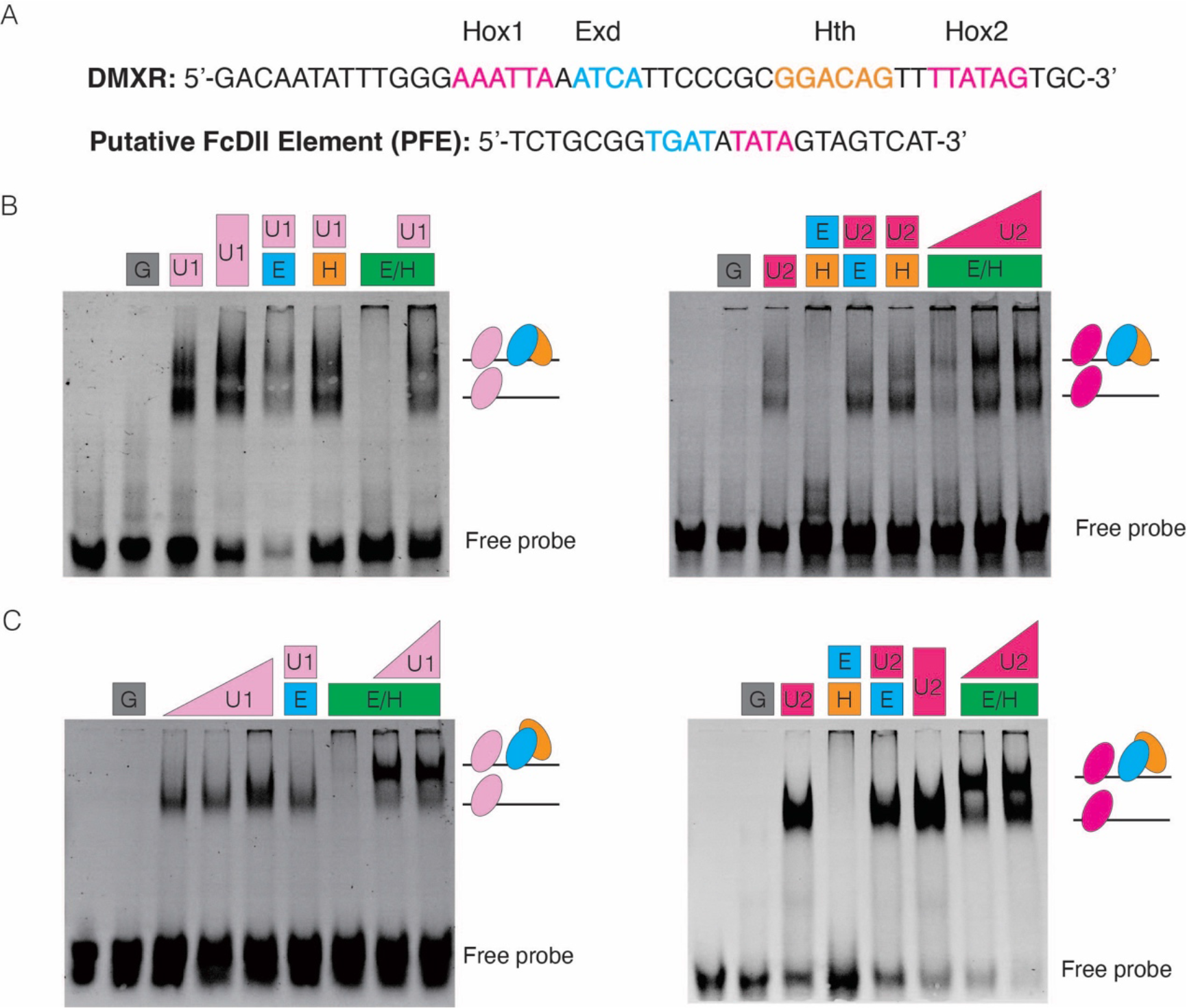
Collembolan Ubx and its cofactors could bind to the DNA elements that contain Hox/Exd/Hth binding motifs. **A.** DNA probes used for EMSAs. DMXR, the transcription regulatory element of *D. melanogaster Dll* (Gebelein et al., 2002; Gebelein et al., 2004); Putative FcDll Element (PFE), a screened DNA region of *F*. *candida*, containing the Hox and Exd binding motifs. The binding sites of Hox/Exd/Hth are shown in colours. **B, C.** Assemblies of collembolan Ubx/Exd/Hth on DMXR and Putative FcDll Element (PFE), respectively. G, GST; U1, FcU1; U2, FcU2; E, Exd; H, Hth. Simplified complexes are indicated on the right.

To test the binding capacity of collembolan Ubx/Exd/Hth complex on its *Dll* DNA, we searched for *Dll* regulatory elements within approximately 4000 bp of the intergenic genomic region upstream from the first exon of *Dll* (Supplementary Data 6). We identified a binding motif containing both Hox/Exd binding sites and referred to this element as the Putative FcDll Element (PFE) (Figure 4A, Supplementary Data 6). The EMSAs show that FcUbx1 and FcUbx2 cooperatively bind with Exd and Hth on this DNA element (Figure 4C), providing additional evidence that collembolan Ubx is proficient in binding DNA defined as the endogenous Hox/Exd binding element. Nonetheless, owing to the absence of practical genetics tools in collembolans, the validation of whether this DNA element serves as the regulatory region of *Dll* in collembolans remains unattainable.

We conclude that the collembolan regulatory trimeric complex Ubx/Exd/Hth can effectively bind to the Hox/Exd/Hth DNA binding motifs, and the lack of *Dll* repression of collembolan Ubx does not appear to be related to any deficiencies in DNA binding capacity. This implies that the mechanisms underlying the absence of repression may stem from other aspects of the regulation progress.

### The C-terminus of FcUbx contains both repression and regulatory domains

Next, to evaluate the transcriptional activity of Collembolan Ubx on *Dll*, we carried out dual luciferase assays in *Drosophila* S2 cells. This involved assessing the repression capacity of various proteins (Figure 5A, Supplementary Data 5), including the complete, truncated, and chimeric forms of both collembolan and *Drosophila* Ubx proteins on the expression of the firefly luciferase reporter gene under the regulation of the DMXR element (Figure 4A, 5A, Supplementary Data 7).

**Figure 5.**
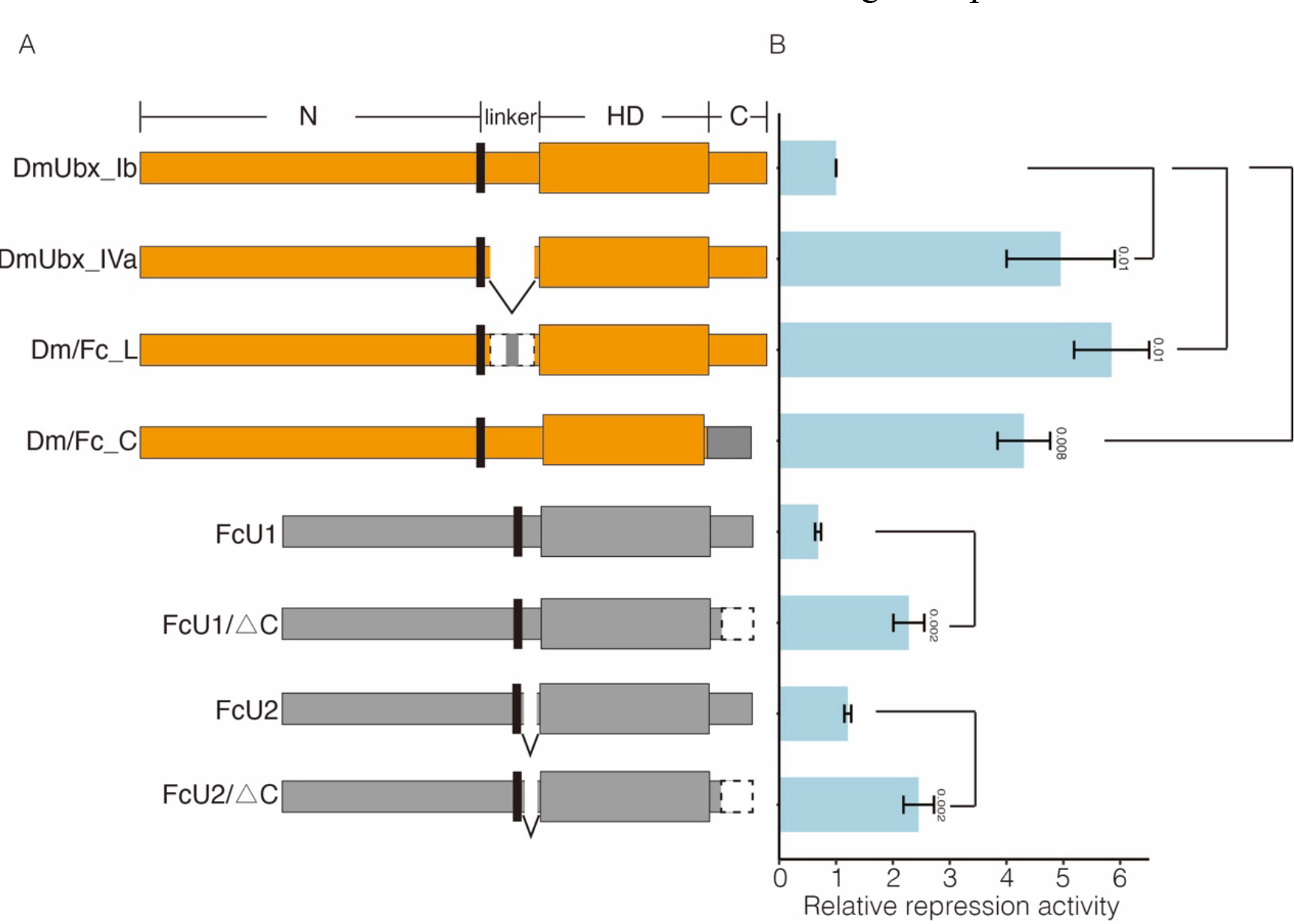
Collembolan Ubx could repress *Dll* transcription while its C-terminus contains regulatory domain(s). **A.** Constructs of proteins used for transcription repression assays, including complete sequences, chimaeras, and truncations of *D. melanogaster* (yellow shade) and *F. candida* (grey shade) Ubx. Dm/Fc_L, the linker of DmUbx_Ib was replaced by the linker (GQSYL) of FcU; Dm/Fc_C, the C-terminus of DmUbx_Ib was replaced by the C-terminus (AKADCKSVY) of FcU; FcU1△C and FcU2△C, the C-terminus (QAQA AKADCKSVY) of FcUbx were deleted. **B.** The relative transcriptional repressive activity of Ubx. The number on the Y-axis indicates transcriptional repression activity relative to DmUbx_Ib. Significant P values of selected pairwise comparisons are shown (Student’s t-test).

Although previous research reported that collembolan Ubx cannot repress *Dll* (Palopoli & Patel, 1998), surprisingly, our results revealed that the two isoforms of collembolan Ubx (FcU1 and FcU2) were capable of inhibiting the expression of the firefly luciferase reporter gene (Figure 5B), which could support our hypothesis that Ubx could repress *Dll* expression during the stages of appendage maturation (Figure 3G). For collembolan Ubx, the linker region is not required for the repression function; In contrast, the longest linker is indispensable for the repression function of *Drosophila* Ubx, as the non-linker isoform of *Drosophila* Ubx (DmUbx_IVa) and the chimeric protein Dm/Fc_L (Linker of *Drosophila* Ubx is replaced by a short linker GQSYL from *F. candida*) significantly lost its repression capability (Figure 5B).

Gebelein et al. revealed that the truncated form of the *Drosophila* Ubx C-terminus is partially able to repress *Dll* (Gebelein et al., 2002). In contrast, our repression assay showed that the truncation of FcU1 and FcU2, which lack a C-terminus (FcU1△C and FcU2△C), significantly decreased their repression ability (Figure 5B), indicating that the C-terminus may contain a potential transcriptional inhibitory domain. Unexpectedly, the chimeric protein Dm/Fc_C (C-terminus of *Drosophila* Ubx is replaced by a short C-terminus AKADCKSVY from *F. candida*) exhibited a substantial loss of repression capacity when compared to DmUbx_Ib, suggesting that the C-terminus of collembolan Ubx (AKADCKSVY) probably contains potential regulatory or modification site(s) that are capable of regulating the repression function of DmUbx_Ib (Figure 5, 6B).

Collectively, our results indicate that the linker of FcUbx is unnecessary for the repression function, while the C-terminus may exhibit both a repression domain (QAQA domain) and a functional regulatory site (S) (Figure 6). These findings provide functional evidence on the mechanisms of Ubx-mediated gene repression.

**Figure 6.**
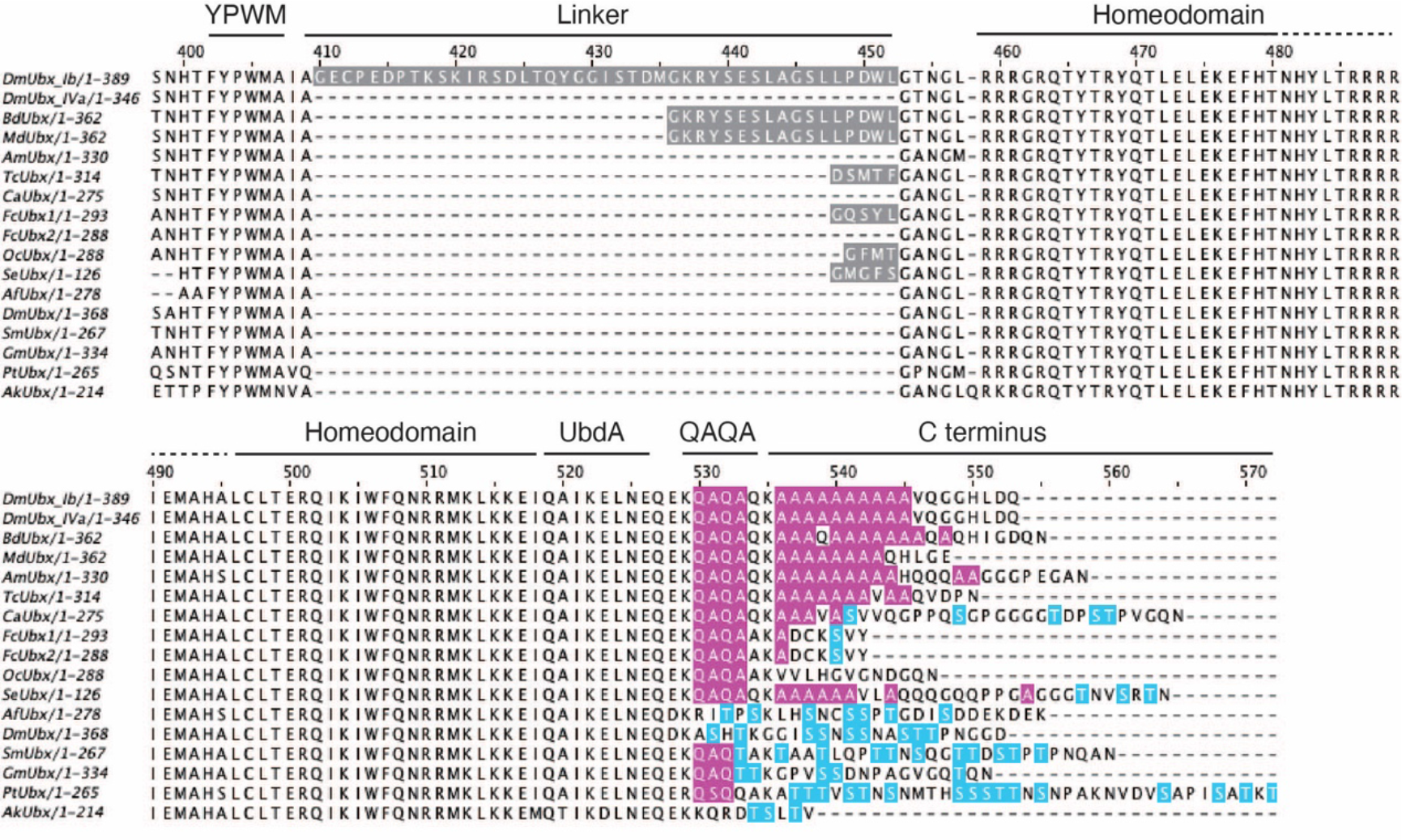
Sequence comparison of the linker and C-terminus of the panarthropod Ubx Partial multiple sequence alignment of Ubx across Panarthropoda, encompassing Insecta, Diplura, Collembola, Protura, Crustacea, Myriapoda, Chelicerata and Onychophora. Insecta: *Drosophila melanogaster* (DmUbx_Ib, DmUbx_IVa), *Bactrocera dorsalis* (BdUbx), *Musca domestica* (MdUbx), *Apis mellifera* (AmUbx), *Tribolium castaneum* (TcUbx); Diplura: *Campodea augens* (CaUbx); Collembola: *Folsomia candida* (FcUbx1, FcUbx2), *Orchesella cincta* (OcUbx); Protura: *Sinentomon erythranum* (SeUbx); Crustacea: *Artemia franciscana* (AfUbx), *Daphnia magna* (DmUbx); Myriapoda: *Strigamia maritima* (SmUbx), *Glomeris marginata* (GmUbx); Chelicerata: *Parasteatoda tepidariorum* (PtUbx); Onychophora: *Acanthokara kaputensis* (AkUbx). The functional domains are indicated. YPWM, YPWM motif; UbdA, UbdA motif. Features of sequences are highlighted in colours. The QAQA domain and poly-Ala stretch are marked in pink, and putative phosphorylated sites S/T (Ser/Thr) are marked in blue. Linkers are marked in grey.

### The sequence evolution of linker and C-terminus in arthropod Ubx

The regulatory functions of the linker and C-terminus of Ubx in *D. melanogaster* and *F. candida* suggest their significant roles in the evolution and regulation of arthropod abdominal appendage formation. To depict the evolutionary trajectory of those sequence features, we compared Ubx sequences from diverse representatives of panarthropods, including Insecta, Diplura, Collembola, Protura, Crustacea, Myriapoda, Chelicerate, and Onychophoran (Supplementary Data 4).

As illustrated in the multiple sequence alignment of Ubx across panarthropods (Figure 6), *D. melanogaster* (DmUbx_Ib) exhibits the longest linker region, consisting of more than 40 amino acids. Nevertheless, the Ubx from proturan (SeUbx), and collembolans (FcUbx1, OcUbx) display markedly shorter linker regions, typically composed of only 3-5 amino acids. In certain species, this linker region may be absent. Remarkably, the sequences of the C-terminus display evolutionary patterns. In Crustaceans, their Ubx (AfUbx, DmUbx) consistently maintain potential Ser/Thr (S/T) phosphorylation sites (Ronshaugen et al., 2002) (Figure 6). In contrast, the insects, characterized by the loss of abdominal appendages in adult stages, feature a QAQA domain and poly-Ala stretches instead of phosphorylation sites (Galant & Carroll, 2002; Ronshaugen et al., 2002) in their Ubx (DmUbx_Ib, DmUbx_IVa, DdUbx, MdUbx, AmUbx and TcUbx) (Figure 6). Interestingly, in the three groups of basal hexapods, proturans maintain short, cylindrical appendages on the first three abdominal segments (Headrick & Gordh, 2009), diplurans possess a pair of cerci at the last abdominal segments (Headrick & Gordh, 2009), and their Ubx (proturan SeUbx and dipluran CaUbx) have QAQA domains, poly-Ala stretches and S/T sites. Meanwhile, in Collembola, which bears three types of abdominal appendages (Figure 1C), the collembolans Ubx (FcUbx and OcUbx) lack poly-Ala stretches but contain the QAQA domains and a putative phosphorylated site Ser (S) in FcUbx (Figure 6).

Genes with similar expression patterns usually participate in the same biological process (Ala et al., 2008; Bar-Joseph, 2004; Bhar et al., 2013; van Dam et al., 2018).

To infer the potential regulation of collembolan Ubx, we conducted Spearman correlation (r > 0.95, *p* < 0.01) analysis throughout embryogenesis (E_0.5d to E_9.5d) and identified a total of 113 genes with 287 transcripts coexpressed with *Ubx*, including the Hox gene *AbdA*. We also found phosphorylation kinase receptors (*ROR*) and phosphatases (*Ptp69D*, *Ptp99A*) (Supplementary Data 8). Given the sequence comparison and functions of FcUbx, we hypothesize that collembolan Ubx may undergo direct or indirect regulation by protein phosphorylation and dephosphorylation during embryogenesis.

## Discussion

### Genes involved in the development of collembolan appendages

Collembola, as a basal hexapod group, exhibits morphological traits that intermediate between crustaceans and insects (Gao et al., 2008; Luan et al., 2005; Misof et al., 2014). Exploring the evolutionary and developmental mechanisms underlying these phenotypes could provide valuable insights into the terrestrialization of hexapods (van Straalen, 2021). In this study, we employed two distinct data mining approaches to comprehensively identify genes involved in collembolan appendage development.

The first approach, the “transcriptome-wide screening strategy,” involves selecting specific appendage formation stages to identify genes that follow the predefined developmental trajectory. We identified several relevant appendage formation genes, such as *Notch*, *En*, *Scr*, *Antp*, *Ubx*, *Exd*, *Hth* and *Lim1* (Table 1), which have been extensively studied in *D. melanogaster* for decades. Engrailed controls the segmentation of embryo (van de Heuvel et al., 1993) and Notch regulates the segmentation of leg (de Celis et al., 1998; Rauskolb & Irvine, 1999); Hox genes Scr, Antp, Ubx are required for the thoracic segment identity (Kaufman & Abbott, 1984), Antp controls the formation of leg (Struhl, 1982) and Ubx repress the abdomen leg (Castelli-Gair & Akam, 1995; Gebelein et al., 2002). Exd, Hth and Lim1 are essential for establishing the proximal-distal axis of limbs and regulating the formation of the coxa, femur, tibia and tarsus (Costello et al., 2015; Ruiz-Losada et al., 2018; Tsuji et al., 2000).

The second approach, the “candidate-gene focusing strategy”, is designed by focusing on the pivotal *Ubx* gene and extracting coexpressed genes during embryogenesis. This analysis not only uncovered the canonical *AbdA* genes (Averof & Patel, 1997; Casares et al., 1996; Konopova & Akam, 2014) but also identified genes related to appendage formation (Supplementary Data 8), including *Lim1* and *DAAM.* In *D. melanogaster*, the deletion of DAAM results in abnormalities in actin filament structures (Barkó et al., 2010; Prokop et al., 2011).

These two synergistic approaches have elucidated a set of genes that have not undergone extensive investigation in arthropods except for *D. melanogaster*. Our analysis introduces new perspectives and broadens the scope of research directions. However, it is crucial to note that the transcriptomes we examined were derived from whole animals, and the identified genes might also be involved in various biological processes, such as organogenesis and neurogenesis. Further research studies are necessary to verify the functional roles of these genes during collembolan appendage development.

### The evolution of functional domains in arthropod Ubx

Throughout its evolution, arthropod Ubx progressively acquired the capacity to inhibit *Dll* expression in the abdomen of insect adults, consequently resulting in the loss of abdominal appendages (Jockusch & Smith, 2015). In this study, by integrating functional assays in collembolan Ubx, we reconstructed the evolutionary trajectory of functional domains in Ubx (Figure 7, Table 2): (1) The ancestral Ubx in Panarthropoda (onychophorans and arthropods) exhibited a consistent DNA binding capability (Galant & Carroll, 2002; Gebelein et al., 2004; Ronshaugen et al., 2002). However, their effectiveness in inhibiting the downstream target *Dll* varied among species (Table 2) (Galant & Carroll, 2002; Gebelein et al., 2002; Grenier & Carroll, 2000; Ronshaugen et al., 2002). (2) In Crustacea, the crustacean Ubx (AfUbx) is unable to inhibit *Dll*, and previous research has proposed that potential regulatory phosphorylation sites may be located in the C-terminus of AfUbx (Galant & Carroll, 2002; Ronshaugen et al., 2002). (3) In Hexapoda, *Drosophila* Ubx (DmUbx_Ib) robustly represses *Dll* expression, the linker region and the C-terminus (including both the QAQA domain and the poly-Ala stretch) play crucial roles in this regulatory process (Galant & Carroll, 2002; Gebelein et al., 2002; Ronshaugen et al., 2002). (4) Remarkably, collembolan Ubx demonstrated the ability to inhibit *Drosophila Dll* expression, mirroring the capability observed in DmUbx_Ib (Figure 5). Nonetheless, (5) unlike *Drosophila* Ubx, the linker region of collembolan Ubx was not essential for this repression (Figure 6). Rather, (6) it appeared that the functional repression domain and potential regulatory phosphorylation sites may be in the C-terminus of collembolan Ubx (Figure 5, 6).

**Figure 7.**
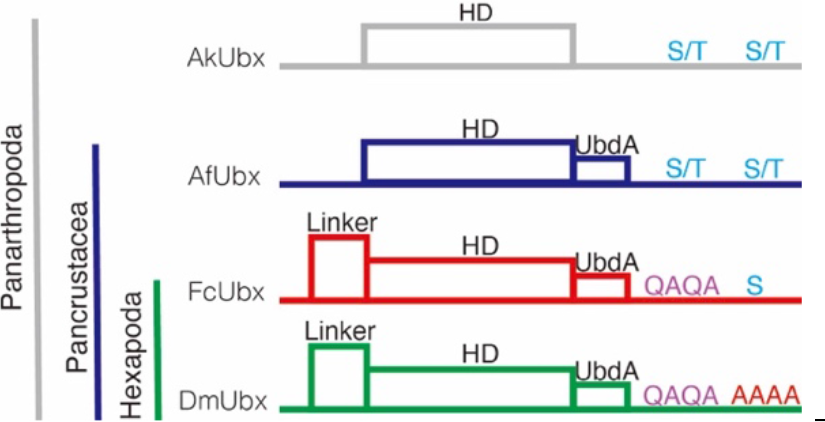
Schematic summary of the functional domains in Ubx of panarthropods. **Note**: +: strong phenotype; -: no phenotype; ND: no data

**Table 2.**
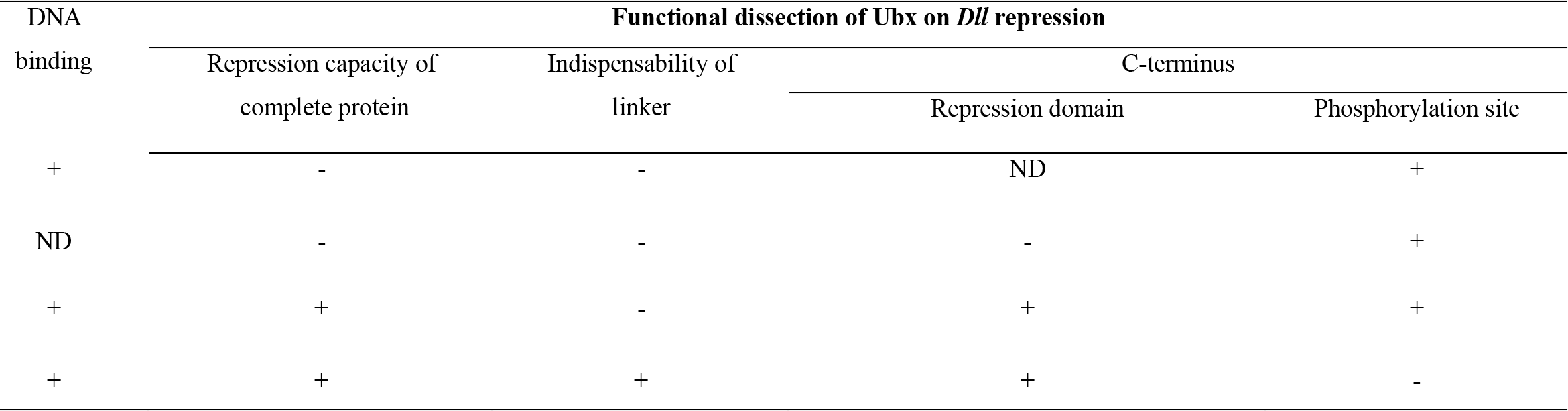
Functional evolution of Ubx.

On the basis of these results, we summarize and propose a functional mechanism of how the collembolan Ubx suppresses *Dll* transcription that appears to be intermediate between that of crustaceans and insects (Table 2). Moreover, given the evidence of characters (QAQA domain, poly Ala stretch and S/T site) in the C-terminus of dipluran and proturan Ubx (Figure 6), we speculate that the ancestral hexapod Ubx could bind and repress the expression of *Dll*. However, the scarcity of Ubx sequences in basal hexapods limits our understanding. Currently, only two complete collembolan Ubx sequences are publicly available (FcUbx and OcUbx), exhibiting inconsistencies in sequence features (Figure 6). With the advancement of genomic sequencing techniques, we anticipate that more Ubx sequences could be extracted from the genomes of basal hexapods. This, in turn, would facilitate the consolidation of sequence features in hexapods and enlighten the exploration of the evolutionary trajectory of functional domains in arthropod Ubx.

### The regulatory mechanism of collembolan Ubx on *Dll*

It is essential to highlight that the discussion on the functional evolution of Ubx primarily relies on the mechanism through which arthropod Ubx suppresses *Drosophila Dll*, and the *bona fide* mechanism of how arthropod Ubx (*trans-factor*) interacts with endogenous regulatory element of *Dll* (*cis-element*) remains underexplored. In combination with our functional assays and transcriptomic analyses, we propose a model to discuss the potential mechanism of how collembolan Ubx regulates the abdominal segments: collembolan Ubx exhibits binding and repression of *Dll* expression uniformly across all segments at the abdominal appendage formation stage (Figures 3F, 4C, 5). Nevertheless, within each of the abdominal segments, collembolan Ubx implements its repression function in a spatially and temporally context-specific manner as depicted in our model (Figure 8). During the early abdominal appendage formation stage (E_3.5d) in abdominal segments 1 and 3 (A1 and A3), Ubx is likely subject to protein phosphorylation, potentially related to Ror (Table 1, Supplementary Data 2, 8). This phosphorylation impairs repression on *Dll*, thereby promoting the development of the ventral tube and retinaculum. We propose that distinct morphologies of these abdominal appendages may be influenced by different cofactors, potentially contributing to morphological malformations, such as Lim1 and CG7526 (Table 1, Supplementary Data 8). We hypothesis two scenarios for abdominal segment 2 (A2) or the appendage maturation stages (E_5.5d onwards): (H1) Ubx undergoes dephosphorylation by phosphatases Ptp69D or Ptp99A (Supplementary Data 8), allowing it to exert its repression function on *Dll*, resulting in the absence of appendages in A2; or (H2) it is still possible that the chromatin landscape of the *Dll* regulatory region becomes inaccessible, thus preventing *Dll* transcription and appendage formation.

**Figure 8.**
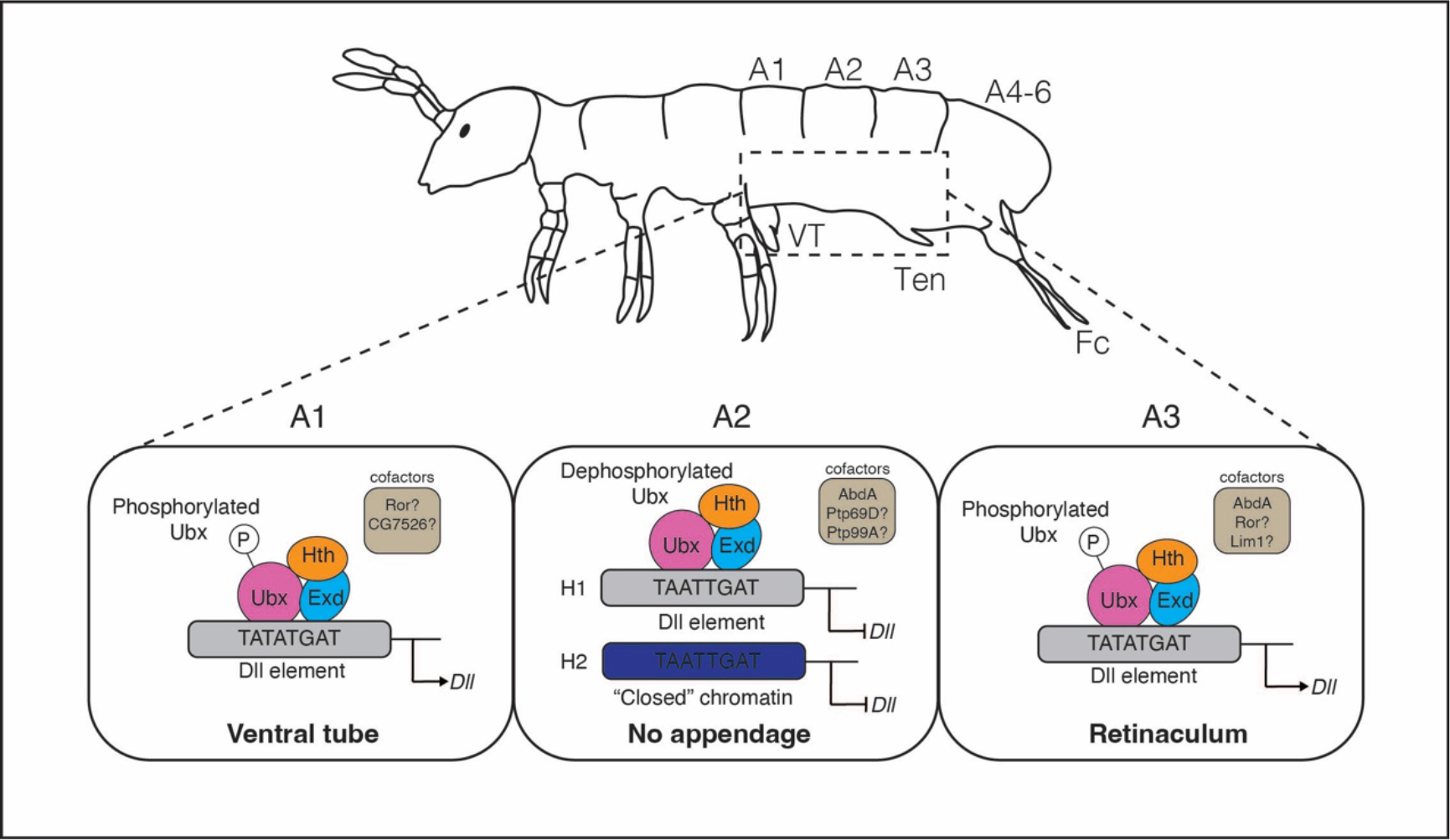
The proposed model for the regulation of collembolan Ubx on *Dll* A schematic morphology of adult *F. candida*. The abdominal segments are shown as A1 to A6. VT, ventral tube; Ten, retinaculum; Fc, furca. The functional regulatory model of Ubx: in A1 and 3, Ubx is phosphorylated, resulting in the derepression of *Dll* and facilitating the formation of appendages; Conversely, in A2, (H1) Ubx exerts its repression function on *Dll* expression; or (H2) the chromatin of *Dll* regulatory region is “closed”, therefore suppressing appendage formation. See the text in the discussion for a detailed explanation.

Despite the current scarcity of genetic tools in collembolans, the confirmation of our model demands leveraging cutting-edge experiments and functional genomics techniques. For instance, ChIP-seq (Park, 2009) or CUT& TAG (Kaya-Okur et al., 2019) allow the identification of Ubx-*Dll* interaction in a genome-wide scale. Additionally, employing the CRISPR/CAS technology (Akinci et al., 2021) to manipulate the Putative FcDll Element would validate its role as a transcriptional regulatory element in collembolans. Furthermore, the incorporation of spatial transcriptomics (Marx, 2021) and spatial ATAC-seq (Deng et al., 2022) to comprehensively map the intricate context of transcription factors and the regulatory landscape in each segment. These methodologies will empower us to investigate how Ubx works precisely, provide compelling evidence for holistic reconstruction of its functional evolution, and ultimately shed light on how arthropods lost their abdominal appendages during evolution.

## Abbreviation

Ubx: Ultrabithorax
AbdA: Abdominal A
Exd: Extradentical
Hth: Homothorax
Dll: Distal-less
Vt: Ventral tube
Ten: Retinaculum
Fc: Furca
An: antenna
AnS: antenna segment
Cllr: clypeolabrum
DO: dorsal organ
IcS: intercalary segment
MdS: mandibular segment
SEM: Scanning electron microscopy
STEM: Short Time-series Expression Miner
RPKM: reads per kilobase per million
GO term: Gene Ontology term
KEGG pathway: Kyoto Encyclopedia of Genes and Genomes
IPTG: isopropyl β-D-1-thiogalactopyranoside
GST: Glutathione S-transferase
DMXR: the transcription regulatory element of D. melanogaster Dll304
PFE: Putative FcDll Element
EMSA: Electrophoretic mobility shift assay

## Supplementary Data

1. The gene expression dynamics of *F. candida*
2. Short Time-series Expression Miner analysis
3. The GO annotation and KEGG pathways of Cluster 42
4. Multiple sequence alignment of Ubx
5. Truncated proteins of Ubx/Exd/Hth
6. Genomic sequence of *F. candida Dll*
7. Dual luciferase assays of collembolan and *Drosophila* proteins
8. Genes coexpressed with *FcUbx*

## Authors’ contributions

Y.L. and Y.-X.L. conceived the project. Y.L. performed all the experiments and data analyses and drafted the manuscript. The authors read and approved the manuscript.

## Acknowledgements

We thank all the lab members from Yin and Luan’s previous laboratory for their help and valuable discussion about this work. We would like to express appreciation to many colleagues. Dr. WU Wei and the team from the Core Facility of Drosophila Resource and Technology at the Centre for Excellence in Molecular Cell Science, CAS for their assistance in establishing the transgenic fly strains of collembolan Ubx and providing troubleshooting support. Miss LI Jiqing and Mr GAO Xiaoyan from the Electron Microscope Platform at the Centre for Excellence in Molecular Plant Sciences, CAS for the aid in the SEM of the collembolans. Dr. ZHAO Qin and Dr. WANG Jing for their suggestions on protein expression. Dr. HE Juanmei and Dr. LANG Nannan for generously sharing the EMSA protocol. Dr. JIA Qiangqiang for assistance with luciferase assays in *Drosophila* S2 cells. Dr. ZHANG Yu for help in bioinformatic analysis. Dr. Chema Martin at Queen Mary University of London and Dr. Kamil Jaron at Wellcome Sanger Institute for the comments on the manuscript.

## References

1. Akinci, E., Hamilton, M. C., Khowpinitchai, B., & Sherwood, R. I. (2021). Using CRISPR to understand and manipulate gene regulation. Development, 148(9). 10.1242/dev.182667

2. Ala, U., Piro, R. M., Grassi, E., Damasco, C., Silengo, L., Oti, M., Provero, P., & Di Cunto, F. (2008). Prediction of human disease genes by human-mouse conserved coexpression analysis. PLoS Comput Biol, 4(3), e1000043. 10.1371/journal.pcbi.1000043

3. Altschul, S. F., Gish, W., Miller, W., Myers, E. W., & Lipman, D. J. (1990). Basic local alignment search tool. J Mol Biol, 215(3), 403–410. 10.1016/s0022-2836(05)80360-2

4. Angelini, & Kaufman. (2005a). Comparative Developmental Genetics and the Evolution of Arthropod Body Plans. Annual Review of Genetics, 39(1), 95–119. 10.1146/annurev.genet.39.073003.112310

5. Angelini, & Kaufman. (2005b). Insect appendages and comparative ontogenetics. Dev Biol, 286(1), 57–77. 10.1016/j.ydbio.2005.07.006

6. Averof, M., & Patel, N. H. (1997). Crustacean appendage evolution associated with changes in Hox gene expression. Nature, 388(6643), 682–686. 10.1038/41786

7. Bairoch, A., Apweiler, R., Wu, C. H., Barker, W. C., Boeckmann, B., Ferro, S., Gasteiger, E., Huang, H., Lopez, R., Magrane, M., Martin, M. J., Natale, D. A., O’Donovan, C., Redaschi, N., & Yeh, L. S. (2005). The Universal Protein Resource (UniProt). Nucleic Acids Res, 33(Database issue), D154-159. 10.1093/nar/gki070

8. Bar-Joseph, Z. (2004). Analyzing time series gene expression data. Bioinformatics, 20(16), 2493–2503. 10.1093/bioinformatics/bth283

9. Bar-Joseph, Z., Gitter, A., & Simon, I. (2012). Studying and modelling dynamic biological processes using time-series gene expression data. Nature Reviews Genetics, 13(8), 552–564. 10.1038/nrg3244

10. Barkó, S., Bugyi, B., Carlier, M. F., Gombos, R., Matusek, T., Mihály, J., & Nyitrai, M. (2010). Characterization of the biochemical properties and biological function of the formin homology domains of Drosophila DAAM. J Biol Chem, 285(17), 13154–13169. 10.1074/jbc.M109.093914

11. Berger, M. F., Badis, G., Gehrke, A. R., Talukder, S., Philippakis, A. A., Peña-Castillo, L., Alleyne, T. M., Mnaimneh, S., Botvinnik, O. B., Chan, E. T., Khalid, F., Zhang, W., Newburger, D., Jaeger, S. A., Morris, Q. D., Bulyk, M. L., & Hughes, T. R. (2008). Variation in homeodomain DNA binding revealed by high-resolution analysis of sequence preferences. Cell, 133(7), 1266–1276. 10.1016/j.cell.2008.05.024

12. Bhar, A., Haubrock, M., Mukhopadhyay, A., Maulik, U., Bandyopadhyay, S., & Wingender, E. (2013). Coexpression and coregulation analysis of time-series gene expression data in estrogen-induced breast cancer cell. Algorithms Mol Biol, 8(1), 9. 10.1186/1748-7188-8-9

13. Brower, D. L. (1986). Engrailed gene expression in Drosophila imaginal discs. Embo j, 5(10), 2649–2656. 10.1002/j.1460-2075.1986.tb04547.x

14. Brown, J. B., Boley, N., Eisman, R., May, G. E., Stoiber, M. H., Duff, M. O., Booth, B. W., Wen, J., Park, S., Suzuki, A. M., Wan, K. H., Yu, C., Zhang, D., Carlson, J. W., Cherbas, L., Eads, B. D., Miller, D., Mockaitis, K., Roberts, J., Davis, C. A., Frise, E., Hammonds, A. S., Olson, S., Shenker, S., Sturgill, D., Samsonova, A. A., Weiszmann, R., Robinson, G., Hernandez, J., Andrews, J., Bickel, P. J., Carninci, P., Cherbas, P., Gingeras, T. R., Hoskins, R. A., Kaufman, T. C., Lai, E. C., Oliver, B., Perrimon, N., Graveley, B. R., & Celniker, S. E. (2014). Diversity and dynamics of the Drosophila transcriptome. Nature, 512(7515), 393–399. 10.1038/nature12962

15. Browne, W. E., & Patel, N. H. (2000). Molecular genetics of crustacean feeding appendage development and diversification. Seminars in Cell & Developmental Biology, 11(6), 427–435. 10.1006/scdb.2000.0196

16. Budd, G. E., & Telford, M. J. (2009). The origin and evolution of arthropods. Nature, 457(7231), 812–817. 10.1038/nature07890

17. Buffry, A. D., Kittelmann, S., & McGregor, A. P. (2023). Characterisation of the role and regulation of Ultrabithorax in sculpting fine-scale leg morphology [Original Research]. Frontiers in Cell and Developmental Biology, 11. 10.3389/fcell.2023.1119221

18. Casares, F., Calleja, M., & Sánchez-Herrero, E. (1996). Functional similarity in appendage specification by the Ultrabithorax and abdominal-A Drosophila HOX genes. Embo j, 15(15), 3934–3942. https://www.ncbi.nlm.nih.gov/pmc/articles/PMC452108/pdf/emboj00015-0170.pdf

19. Castelli-Gair, J., & Akam, M. (1995). How the Hox gene Ultrabithorax specifies two different segments: the significance of spatial and temporal regulation within metameres. Development, 121(9), 2973–2982. 10.1242/dev.121.9.2973

20. Cohen, B., Simcox, A. A., & Cohen, S. M. (1993). Allocation of the thoracic imaginal primordia in the Drosophila embryo. Development, 117(2), 597–608. 10.1242/dev.117.2.597

21. Cohen, B., Wimmer, E. A., & Cohen, S. M. (1991). Early development of leg and wing primordia in the Drosophila embryo. Mech Dev, 33(3), 229–240. 10.1016/0925-4773(91)90030-a

22. Cohen, S. M. (1990). Specification of limb development in the Drosophila embryo by positional cues from segmentation genes. Nature, 343(6254), 173–177. 10.1038/343173a0

23. Conesa, A., Götz, S., García-Gómez, J. M., Terol, J., Talón, M., & Robles, M. (2005). Blast2GO: a universal tool for annotation, visualization and analysis in functional genomics research. Bioinformatics, 21(18), 3674–3676. 10.1093/bioinformatics/bti610

24. Costello, I., Nowotschin, S., Sun, X., Mould, A. W., Hadjantonakis, A. K., Bikoff, E. K., & Robertson, E. J. (2015). Lhx1 functions together with Otx2, Foxa2, and Ldb1 to govern anterior mesendoderm, node, and midline development. Genes Dev, 29(20), 2108-2122. 10.1101/gad.268979.115

25. de Celis, J. F., Tyler, D. M., de Celis, J., & Bray, S. J. (1998). Notch signalling mediates segmentation of the Drosophila leg. Development, 125(23), 4617–4626. 10.1242/dev.125.23.4617

26. Deng, Y., Bartosovic, M., Ma, S., Zhang, D., Kukanja, P., Xiao, Y., Su, G., Liu, Y., Qin, X., Rosoklija, G. B., Dwork, A. J., Mann, J. J., Xu, M. L., Halene, S., Craft, J. E., Leong, K. W., Boldrini, M., Castelo-Branco, G., & Fan, R. (2022). Spatial profiling of chromatin accessibility in mouse and human tissues. Nature, 609(7926), 375–383. 10.1038/s41586-022-05094-1

27. Durston, A. J., Jansen, H. J., In der Rieden, P., & Hooiveld, M. H. (2011). Hox collinearity - a new perspective. Int J Dev Biol, 55(10-12), 899–908. 10.1387/ijdb.113358ad

28. Ekker, S. C., Young, K. E., von Kessler, D. P., & Beachy, P. A. (1991). Optimal DNA sequence recognition by the Ultrabithorax homeodomain of Drosophila. Embo j, 10(5), 1179–1186. 10.1002/j.1460-2075.1991.tb08058.x

29. Ernst, J., & Bar-Joseph, Z. (2006). STEM: a tool for the analysis of short time series gene expression data. BMC Bioinformatics, 7, 191. 10.1186/1471-2105-7-191

30. Ernst, J., Nau, G. J., & Bar-Joseph, Z. (2005). Clustering short time series gene expression data. Bioinformatics, 21 *Suppl 1*, i159–168. 10.1093/bioinformatics/bti1022

31. Faddeeva-Vakhrusheva, A., Kraaijeveld, K., Derks, M. F. L., Anvar, S. Y., Agamennone, V., Suring, W., Kampfraath, A. A., Ellers, J., Le Ngoc, G., van Gestel, C. A. M., Mariën, J., Smit, S., van Straalen, N. M., & Roelofs, D. (2017). Coping with living in the soil: the genome of the parthenogenetic springtail Folsomia candida. BMC Genomics, 18(1), 493. 10.1186/s12864-017-3852-x

32. Fountain, M. T., & Hopkin, S. P. (2005). Folsomia candida (Collembola): a “standard” soil arthropod. Annu Rev Entomol, 50(Volume 50, 2005), 201-222. 10.1146/annurev.ento.50.071803.130331

33. Fountain, M. T., & Hopkin, S. P. (2005). FOLSOMIA CANDIDA (COLLEMBOLA): A “Standard” Soil Arthropod. Annual Review of Entomology, 50(1), 201–222. 10.1146/annurev.ento.50.071803.130331

34. Galant, R., & Carroll, S. B. (2002). Evolution of a transcriptional repression domain in an insect Hox protein. Nature, 415(6874), 910–913. 10.1038/nature717

35. Gao, Y., Bu, Y., Luan, Y.-X., & Yin, W.-Y. (2006). Preliminary Observation on the Embryonic Development of Folsomia candida (Collembola: Isotomidae). Zoological Research, 5(27), 519–524.

36. Gao, Y., Bu, Y., & Luan, Y. X. (2008). Phylogenetic relationships of basal hexapods reconstructed from nearly complete 18S and 28S rRNA gene sequences. Zoolog Sci, 25(11), 1139–1145. 10.2108/zsj.25.1139

37. Gaunt, S. J. (2015). The significance of Hox gene collinearity. Int J Dev Biol, 59(4-6), 159–170. 10.1387/ijdb.150223sg

38. Gebelein, B., Culi, J., Ryoo, H. D., Zhang, W., & Mann, R. S. (2002). Specificity of Distalless repression and limb primordia development by abdominal Hox proteins. Dev Cell, 3(4), 487–498. 10.1016/s1534-5807(02)00257-5

39. Gebelein, B., McKay, D. J., & Mann, R. S. (2004). Direct integration of Hox and segmentation gene inputs during Drosophila development. Nature, 431(7009), 653–659. 10.1038/nature02946

40. Geyer, A., Koltsaki, I., Hessinger, C., Renner, S., & Rogulja-Ortmann, A. (2015). Impact of Ultrabithorax alternative splicing on Drosophila embryonic nervous system development. Mechanisms of Development, 138, 177–189. 10.1016/j.mod.2015.08.007

41. Giribet, G., Edgecombe, G. D., & Wheeler, W. C. (2001). Arthropod phylogeny based on eight molecular loci and morphology. Nature, 413(6852), 157–161. 10.1038/35093097

42. Grenier, J. K., & Carroll, S. B. (2000). Functional evolution of the Ultrabithorax protein. Proc Natl Acad Sci U S A, 97(2), 704–709. 10.1073/pnas.97.2.704

43. He, J. M., Zhu, H., Zheng, G. S., Liu, P. P., Wang, J., Zhao, G. P., Zhu, G. Q., Jiang, W. H., & Lu, Y. H. (2016). Direct Involvement of the Master Nitrogen Metabolism Regulator GlnR in Antibiotic Biosynthesis in Streptomyces. J Biol Chem, 291(51), 26443–26454. 10.1074/jbc.M116.762476

44. Headrick, D. H., & Gordh, G. (2009). Chapter 5 - Anatomy: Head, Thorax, Abdomen, and Genitalia. In V. H. Resh & R. T. Cardé (Eds.), Encyclopedia of Insects *(Second Edition)* (pp. 11-21). Academic Press. 10.1016/B978-0-12-374144-8.00005-9

45. Hughes, & Kaufman. (2002a). Exploring the myriapod body plan: expression patterns of the ten Hox genes in a centipede. Development, 129(5), 1225–1238. 10.1242/dev.129.5.1225

46. Hughes, & Kaufman. (2002b). Hox genes and the evolution of the arthropod body plan. Evol Dev, 4(6), 459–499. 10.1046/j.1525-142x.2002.02034.x

47. Jockusch, E. L., & Smith, F. W. (2015). Hexapoda: Comparative Aspects of Later Embryogenesis and Metamorphosis. In A. Wanninger (Ed.), Evolutionary Developmental Biology of Invertebrates 5: Ecdysozoa III: Hexapoda (pp. 111- 208). Springer Vienna. 10.1007/978-3-7091-1868-9_3

48. Jockusch, E. L., Williams, T. A., & Nagy, L. M. (2004). The evolution of patterning of serially homologous appendages in insects. Development Genes and Evolution, 214(7), 324–338. 10.1007/s00427-004-0412-6

49. Kanehisa, M., & Goto, S. (2000). KEGG: kyoto encyclopedia of genes and genomes. Nucleic Acids Res, 28(1), 27–30. 10.1093/nar/28.1.27

50. Kaufman, T. C., & Abbott, M. K. (1984). Homoeotic Genes and the Specification of Segmental Identity in the Embryo and Adult Thorax of Drosophila Melanogaster. In G. M. Malacinski & W. H. Klein (Eds.), Molecular Aspects of Early Development (pp. 189-218). Springer US. 10.1007/978-1-4684-4628-9_9

51. Kaya-Okur, H. S., Wu, S. J., Codomo, C. A., Pledger, E. S., Bryson, T. D., Henikoff, J. G., Ahmad, K., & Henikoff, S. (2019). CUT&Tag for efficient epigenomic profiling of small samples and single cells. Nature Communications, 10(1), 1930. 10.1038/s41467-019-09982-5

52. Kim, D., Pertea, G., Trapnell, C., Pimentel, H., Kelley, R., & Salzberg, S. L. (2013). TopHat2: accurate alignment of transcriptomes in the presence of insertions, deletions and gene fusions. Genome Biol, 14(4), R36. 10.1186/gb-2013-14-4-r36

53. Kolde, R. (2012). Pheatmap: pretty heatmaps. R package version, 1(2), 726.

54. Konopova, B., & Akam, M. (2014). The Hox genes Ultrabithorax and abdominal-A specify three different types of abdominal appendage in the springtail Orchesella cincta (Collembola). Evodevo, 5(1), 2. 10.1186/2041-9139-5-2

55. Krogh, P. H. (2009). Toxicity testing with the collembolans Folsomia fimetaria and Folsomia candida and the results of a ringtest.

56. Kurata, S., Go, M. J., Artavanis-Tsakonas, S., & Gehring, W. J. (2000). Notch signaling and the determination of appendage identity. Proc Natl Acad Sci U S A, 97(5), 2117–2122. 10.1073/pnas.040556497

57. Langmead, B., & Salzberg, S. L. (2012). Fast gapped-read alignment with Bowtie 2. Nat Methods, 9(4), 357–359. 10.1038/nmeth.1923

58. Liang, Y., Xie, W., & Luan, Y. X. (2019). Developmental expression and evolution of hexamerin and haemocyanin from Folsomia candida (Collembola). Insect Molecular Biology, 28(5), 716–727. 10.1111/imb.12585

59. Luan, Y. X., Cui, Y., Chen, W. J., Jin, J. F., Liu, A. M., Huang, C. W., Potapov, M., Bu, Y., Zhan, S., Zhang, F., & Li, S. (2023). High-quality genomes reveal significant genetic divergence and cryptic speciation in the model organism Folsomia candida (collembola). Mol Ecol Resour, 23(1), 273–293. 10.1111/1755-0998.13699

60. Luan, Y. X., Mallatt, J. M., Xie, R. D., Yang, Y. M., & Yin, W. Y. (2005). The phylogenetic positions of three Basal-hexapod groups (protura, diplura, and collembola) based on ribosomal RNA gene sequences. Mol Biol Evol, 22(7), 1579–1592. 10.1093/molbev/msi148

61. Marx, V. (2021). Method of the Year: spatially resolved transcriptomics. Nature Methods, 18(1), 9–14. 10.1038/s41592-020-01033-y

62. Matsuda, R. (2017). Morphology and evolution of the insect abdomen: with special reference to developmental patterns and their bearings upon systematics. Elsevier.

63. McGinnis, S., & Madden, T. L. (2004). BLAST: at the core of a powerful and diverse set of sequence analysis tools. Nucleic Acids Res, 32(Web Server issue), W20- 25. 10.1093/nar/gkh435

64. McIntosh, B. B., & Ostap, E. M. (2016). Myosin-I molecular motors at a glance. J Cell Sci, 129(14), 2689–2695. 10.1242/jcs.186403

65. Mead, T. J., Martin, D. R., Wang, L. W., Cain, S. A., Gulec, C., Cahill, E., Mauch, J., Reinhardt, D., Lo, C., Baldock, C., & Apte, S. S. (2022). Proteolysis of fibrillin- 2 microfibrils is essential for normal skeletal development. eLife, 11, e71142. 10.7554/eLife.71142

66. Misof, B., Liu, S., Meusemann, K., Peters, R. S., Donath, A., Mayer, C., Frandsen, P. B., Ware, J., Flouri, T., Beutel, R. G., Niehuis, O., Petersen, M., Izquierdo- Carrasco, F., Wappler, T., Rust, J., Aberer, A. J., Aspöck, U., Aspöck, H., Bartel, D., Blanke, A., Berger, S., Böhm, A., Buckley, T. R., Calcott, B., Chen, J., Friedrich, F., Fukui, M., Fujita, M., Greve, C., Grobe, P., Gu, S., Huang, Y., Jermiin, L. S., Kawahara, A. Y., Krogmann, L., Kubiak, M., Lanfear, R., Letsch, H., Li, Y., Li, Z., Li, J., Lu, H., Machida, R., Mashimo, Y., Kapli, P., McKenna, D. D., Meng, G., Nakagaki, Y., Navarrete-Heredia, J. L., Ott, M., Ou, Y., Pass, G., Podsiadlowski, L., Pohl, H., von Reumont, B. M., Schütte, K., Sekiya, K., Shimizu, S., Slipinski, A., Stamatakis, A., Song, W., Su, X., Szucsich, N. U., Tan, M., Tan, X., Tang, M., Tang, J., Timelthaler, G., Tomizuka, S., Trautwein, M., Tong, X., Uchifune, T., Walzl, M. G., Wiegmann, B. M., Wilbrandt, J., Wipfler, B., Wong, T. K., Wu, Q., Wu, G., Xie, Y., Yang, S., Yang, Q., Yeates, D. K., Yoshizawa, K., Zhang, Q., Zhang, R., Zhang, W., Zhang, Y., Zhao, J., Zhou, C., Zhou, L., Ziesmann, T., Zou, S., Li, Y., Xu, X., Zhang, Y., Yang, H., Wang, J., Wang, J., Kjer, K. M., & Zhou, X. (2014). Phylogenomics resolves the timing and pattern of insect evolution. Science, 346(6210), 763–767. 10.1126/science.1257570

67. Monteiro, A. S., & Ferrier, D. E. K. (2006). Hox genes are not always Colinear [Review]. International Journal of Biological Sciences, 2(3), 95–103. 10.7150/ijbs.2.95

68. Natori, K., Tajiri, R., Furukawa, S., & Kojima, T. (2012). Progressive tarsal patterning in the Drosophila by temporally dynamic regulation of transcription factor genes. Developmental Biology, 361(2), 450–462. 10.1016/j.ydbio.2011.10.031

69. Noyes, M. B., Christensen, R. G., Wakabayashi, A., Stormo, G. D., Brodsky, M. H., & Wolfe, S. A. (2008). Analysis of homeodomain specificities allows the family- wide prediction of preferred recognition sites. Cell, 133(7), 1277–1289. 10.1016/j.cell.2008.05.023

70. O’Day, K. E. (2006). Notch Signaling and Segmentation in Parhyale Hawaiensis. University of California, Berkeley. https://books.google.co.uk/books?id=FbFPAQAAMAAJ

71. Palopoli, M. F., & Patel, N. H. (1998). Evolution of the interaction between Hox genes and a downstream target. Curr Biol, 8(10), 587–590. 10.1016/s0960-9822(98)70228-3

72. Panganiban, G., Irvine, S. M., Lowe, C., Roehl, H., Corley, L. S., Sherbon, B., Grenier, J. K., Fallon, J. F., Kimble, J., Walker, M., Wray, G. A., Swalla, B. J., Martindale, M. Q., & Carroll, S. B. (1997). The origin and evolution of animal appendages. Proc Natl Acad Sci U S A, 94(10), 5162–5166. 10.1073/pnas.94.10.5162

73. Park, P. J. (2009). ChIP–seq: advantages and challenges of a maturing technology. Nature Reviews Genetics, 10(10), 669–680. 10.1038/nrg2641

74. Passner, J. M., Ryoo, H. D., Shen, L., Mann, R. S., & Aggarwal, A. K. (1999). Structure of a DNA-bound Ultrabithorax-Extradenticle homeodomain complex. Nature, 397(6721), 714-719. 10.1038/17833

75. Peel, A. D., Chipman, A. D., & Akam, M. (2005). Arthropod Segmentation: beyond the Drosophila paradigm. Nature Reviews Genetics, 6(12), 905–916. 10.1038/nrg1724

76. Prokop, A., Sánchez-Soriano, N., Gonçalves-Pimentel, C., Molnár, I., Kalmár, T., & Mihály, J. (2011). DAAM family members leading a novel path into formin research. Commun Integr Biol, 4(5), 538–542. 10.4161/cib.4.5.16511

77. Rauskolb, C., & Irvine, K. D. (1999). Notch-Mediated Segmentation and Growth Control of the Drosophila Leg. Developmental Biology, 210(2), 339–350. 10.1006/dbio.1999.9273

78. Reed, H. C., Hoare, T., Thomsen, S., Weaver, T. A., White, R. A., Akam, M., & Alonso, C. R. (2010). Alternative splicing modulates Ubx protein function in Drosophila melanogaster. Genetics, 184(3), 745–758. 10.1534/genetics.109.112086

79. Ronshaugen, M., McGinnis, N., & McGinnis, W. (2002). Hox protein mutation and macroevolution of the insect body plan. Nature, 415(6874), 914–917. 10.1038/nature716

80. Rozewicki, J., Li, S., Amada, K. M., Standley, D. M., & Katoh, K. (2019). MAFFT- DASH: integrated protein sequence and structural alignment. Nucleic Acids Research, 47(W1), W5–W10. 10.1093/nar/gkz342

81. RStudio. (2020). RStudio: Integrated Development for R. RStudio. http://www.rstudio.com/.

82. Ruiz-Losada, M., Blom-Dahl, D., Córdoba, S., & Estella, C. (2018). Specification and Patterning of Drosophila Appendages. J Dev Biol, 6(3). 10.3390/jdb6030017

83. Ryoo, H. D., Marty, T., Casares, F., Affolter, M., & Mann, R. S. (1999). Regulation of Hox target genes by a DNA bound Homothorax/Hox/Extradenticle complex. Development, 126(22), 5137–5148. 10.1242/dev.126.22.5137

84. Slattery, M., Riley, T., Liu, P., Abe, N., Gomez-Alcala, P., Dror, I., Zhou, T., Rohs, R., Honig, B., Bussemaker, Harmen J., & Mann, Richard S. (2011). Cofactor Binding Evokes Latent Differences in DNA Binding Specificity between Hox Proteins. Cell, 147(6), 1270-1282. 10.1016/j.cell.2011.10.053

85. Struhl, G. (1982). Genes controlling segmental specification in the Drosophila thorax. Proc Natl Acad Sci U S A, 79(23), 7380–7384. 10.1073/pnas.79.23.7380

86. Timmermans, M., Roelofs, D., Mariën, J., & van Straalen, N. M. (2008). Revealing pancrustacean relationships: Phylogenetic analysis of ribosomal protein genes places Collembola (springtails) in a monophyletic Hexapoda and reinforces the discrepancy between mitochondrial and nuclear DNA markers. BMC Evolutionary Biology, 8(1), 83. 10.1186/1471-2148-8-83

87. Trapnell, C., Roberts, A., Goff, L., Pertea, G., Kim, D., Kelley, D. R., Pimentel, H., Salzberg, S. L., Rinn, J. L., & Pachter, L. (2012). Differential gene and transcript expression analysis of RNA-seq experiments with TopHat and Cufflinks. Nat Protoc, 7(3), 562–578. 10.1038/nprot.2012.016

88. Tsuji, T., Sato, A., Hiratani, I., Taira, M., Saigo, K., & Kojima, T. (2000). Requirements of Lim1, a Drosophila LIM-homeobox gene, for normal leg and antennal development. Development, 127(20), 4315–4323. 10.1242/dev.127.20.4315

89. Tully, T., & Potapov, M. (2015). Intraspecific Phenotypic Variation and Morphological Divergence of Strains of Folsomia candida (Willem) (Collembola: Isotomidae), the “Standard” Test Springtaill. PLoS One, 10(9), e0136047. 10.1371/journal.pone.0136047

90. van Dam, S., Võsa, U., van der Graaf, A., Franke, L., & de Magalhães, J. P. (2018). Gene co-expression analysis for functional classification and gene-disease predictions. Brief Bioinform, 19(4), 575–592. 10.1093/bib/bbw139

91. van de Heuvel, M., Klingensmith, J., Perrimon, N., & Nusse, R. (1993). Cell patterning in the Drosophila segment: engrailed and wingless antigen distributions in segment polarity mutant embryos. Development, 119(Supplement), 105–114. 10.1242/dev.119.Supplement.105

92. van Straalen, N. M. (2021). Evolutionary terrestrialization scenarios for soil invertebrates. Pedobiologia, *87-88*, 150753. 10.1016/j.pedobi.2021.150753

93. Waterhouse, A. M., Procter, J. B., Martin, D. M. A., Clamp, M., & Barton, G. J. (2009). Jalview Version 2—a multiple sequence alignment editor and analysis workbench. Bioinformatics, 25(9), 1189-1191. 10.1093/bioinformatics/btp033

94. Weatherbee, S. D., & Carroll, S. B. (1999). Selector Genes and Limb Identity in Arthropods and Vertebrates. Cell, 97(3), 283–286. 10.1016/S0092-8674(00)80737-0

95. Wickham, H. (2016). ggplot2: Elegant Graphics for Data Analysis. https://ggplot2.tidyverse.org

96. Wu, J., & Cohen, S. M. (1999). Proximodistal axis formation in the Drosophila leg: subdivision into proximal and distal domains by Homothorax and Distal-less. Development, 126(1), 109–117. 10.1242/dev.126.1.109

97. Yuzuki, D. (2015). *BGISEQ-500 debuts at the International Congress of Genomics 10*. In *Next Generation Technologist* *[online]* http://www.yuzuki.org/bgiseq-500-debut-at-the-international-congress-of-genomics-10

