## Supplementary material for "The functional evolution of collembolan Ubx on the regulation of abdominal appendage formation": 6-Genomics sequences of F. candida Dll.docx

>dll_combined_gDNA_(F._candida) ATAAAAATAATACAATCGTGACACATTTCATAACTGAAATGTGCGTACAAATAGTTAAACAAAATGGATAGCTGTAGTATTTAAGTACAAGTTTGCTGACAATACCATTTTGAAATTTTAACAGACAATTTTCTACAAATCTCCAGTTTTTAGGTGACAACTGCACTTGTTCACAACAAAGGACACGAATGACAATGGATTCATCGCATCCAACACCGCAAGGACCACCGTGAAATCGGAAATCACTTCACGCCAACTTTGCTCAAGCCTTTCTTTCTCGAGATGACAAAAAAGTGAGCGAGATTCTCAACAAACAATGAGAAATGTGTGTGTTATATATTTGACGAACTTTAATCTTCATTGTGAGACTTTGGCCCAGCTTGGGTCCGTTGTTTTCACATAAAATCGGTTATCGTAACTTTGTATGTTTATGCGAAAGGTGAAATATTGGCTGAAATATAATTCACGAGTTACATATTTATGTAAGTTTTAAGGATTCGTTTGGTCTGGTGCATTCAACTTTAGTGCAGTTGATTATTACTTCATAATTCATCCACTAATTACACGAGGAGGATGTTTGTTCTCATATTAAAACATTATTAAGTTTCAATTACACATACCGTCTTACATTTCGCCAATTACTGTAATAACACACGTGTCCATAAGTACTCAACTTCATACTTCATATATGCATGCATGTATGATGTATGTAAGTACCTCTTGAATTCAATAAAGTTGGAAATATTACAAACATGTACATATCTACATGCACATGTAGCAACTGGCAGAAAATAAGTCGGGTTGAAGTAATCCAAGCATATCTTTCATTAAGTAAAGTTATTGAGTACCCTGGCTTCCGAAACGTGTATGTAGGTGGATAATTGACCATGACGTACGAATACTTTCTCTGCGGTGATATATAGTAGTCATTTTTGAAAATATAATGTACGCACTGTGCCACTCGTCATATTGGGTCCAAGGGAGGAGCAATTATTTTGCGAGTGGCATCCCATCCAACGAAATGCCACCCTTCCAACCGGAAAAGAAAGAAATATGAAATGTGTAGGCTTGCTCAATCAATTCGCGACATGTGGAATTATTAGCATATTTATTAATTAGTGACCACACCCACGCCAAGCTACCCAGGCACAGGAATGACAACATGTACCAGAACTGATCCAGTATGGAAAAGGCAAACAGAGAGAGGGTGACAAGACAAACCCCTCCGGAATACAATACGTGATGTGTACAGACATACACGCAATATGGACGACTTGGTATGTATGTCGACGTGATGAAACAGAAAACTGAAATGTGACAAGAGGATTCTCGATATAGGCACTTGGTGTTGTTTGAAATGTGAGCAATATGGCAAATTAATTAGTTTCAGTCGCCACCATCCACTCAGCCAATCCACTCAGAATGGAGGTCTCCCAACATTTTTCACAGATTATGGCCAAATAATGAATTGATCAGAAATTTAATTAATAACAAATTTGCAAAAGATATTCTCAGCCAATGTGCAGGTTTGGCAGGGATATTTGCATTTGAAGAGGAGAAAACGCACAGTTAATTTTGAAAAGTGTAAAATTTGTCACCATTAGGGCGCGCTTTCTCCTCTTTTGATGCAAATATCTCTGCCAAACCTACGCGTTGAGAGATATTGTCTTTTACAACTTTCATATCACTTAACTTTGTGATCTATTGAGACTAATTTGACCAAAATCGGAGAGCATACGCTGGAAAACCTCCATTTTGAATGCGTAAAACTTTCACCACTAAGCGATTGATTCATCATTGTATGTTATGTAGCTTTTTTCCCTCCACAATAGCAATAATACTAATTCTGAGAGGCTCTTATGCCCATTTCGACTGGGGAGAGGTCGGCTATGATCATAAAAATATGTCATTCGATAATAATGGTAAAAAGACCAATTTATCCATCCACTCACAATAAATCCCGGCTCTTATAACACCAACCAACACCACACGGCCATGAGGAAAGTACAACAACAACAGGACGGGCTGGCATTCGAGGTGTGCCTTTACCCAACACCACAGGCAATTTATCAACTTAACAGGAGACAAGAATAAAAGAGGAAAGCAAGCAGGCAGAAAATTGGGCAATTTAGCGAATTCACCCCAATTCGGGTAATTGAATTGTCTAACAAACACTGACACTACGTAGCAGCCAAAACCCAACCGTTTCAACCTCTCTTGAATTATTTTCTCTACAATCTCGCAGAATTTCCTTCCCCAGAGCAATTTACAGAAATCTAAGAAAAGCAACAGTGGTGCAGTGTCGCGAATTATTTTATTAATAAAAGGACGCCATCGGTGCGTTCAGCAATTCTGCCACAGTTTTCGTGTCAGATCATATTTTATCTCGAGTACTTATTTACCACGACATTACTCACGATTTAATTGATCAAACTTCATTGTCCAACTTTTTCATGGCGTTTATGCATCAACTCCCAACTTTCCCCGCAACTTTCTCCTATTTCGTTGTTGGGAGAAGCTGTTTTTCGAGAAAATTGAGGTCCTAGTGGAATCGTTCCTCTAATATTTTATTCATCCTTCATCTATGTTCTTGGAGAAAGCCACAGGTAAAAGTGGCGGAATAACAGGAGGAGAAGCATGTTTCACCCACCACGGGCCAAACAAGTGCTAAATAAGTTATAAACTAAATTGCACATTTATCCAGCCCAAACTTTCCCGTAAACGGAGCTTATTCATCCATCCCCTCTCATTATTAGGGAAATAATAATTTTATTTTTCTCAATTTAAGAATTTCCCTTCCCTTCATTCGTGTCGACGAGGTGAACATTTATTTCTTTAATTCTTAAATCCCTTCAATTTCCTAATCCATAACATTACTTAAAACGTTTAATTGTGTTTAAATACAAGTCTTTAAATCTCTCAAGGACTGGTCAAATTTCGAGTTTTAGTTTCATTTTAAAATTGCTATGTATGTACCACCAGAAAATCGGCAGGATTATTTGATACACATTTCTAAAAACAAGAGATTTTCTATTTCGACACGATTTTCTGGTTGTGCTCGATTTGATTACAGAATCGAACGATGTAGGCAAGTCTTTGAATAGCGAAATCTTTCGACTCGGAGTCAAAAGTGGTTGGTGGCGTTGGAGGAGGTGGCCTGCGAGGCGTAATCCAGCCCAAGCGTTTCTGAGGCGGAGTCGGAAAACTACCCATCAGCCCAGCACGACCATAGCGTCAGAGCCATAGGAAAAAGGTACCTTTGCTGGGGGTAGTTTAGCATAGCTTTAGTTTAGTATGGGGAAGGTAAGGTTGGATTTGGTTTGGTAAGGTGAGGAATGATTCCCATCCGCAATTACAGTTTCACGAGGGCAAGCGAGCTGACGAGATCACGACTCACCTCACTTATTTACTCCCAACCAACTTCTTGTGAAAAAAGTCTCAATAAATTTATACACGGATCAGCCTCGCTTTTAGATACGATCTGTCATTTGTGGTTTCACAGATAAGCTGCACAAAGATTCCCAGCCCAACGCTTAATGGGCAATTTGTCCACGCTGCTCACTGCTCCTCTTCTTCTAATCCAGACCAATTTGTGGTTTATACGCGTCTCAAATGGTATCGGGAAAAGTGACACAAAACTAGAAACCTAGTTATCTAAAAGATAAGGACACGGAGGAAAAAGATAAGGCTGGAATTTTTTTGGGGGAAATGCAGCATTGAAAAGTGACTGAATTTTACGGAAAGTAAAAATAAAGCCGGCATTCAACAGTTTAGAGTTTTTGTGGTGCACTCATTAAGTGTAATTAGTGAAATTACTCTCACGTGCGCGTACCAACGAACGAATGAGCCCTTCTTCTTCGCCTTACTTCCGCTTGGGTTGAGTGCAGGAAGTGAAAGTTATTTATGAAATAAAACCTCGATTTTGACAAGAATATTGTAAAAGTAGAATTTTTCGAAATTTCAAAGACAACTTGACACGACTTAAGTAAGCTTGATATGGAACTACAGAGTGAGCTGAAACTTTGCAATTTGAATATCCTAGTGACTATTGCGGATAGTGAACATTTGATCAGCTAGAAATAAAGAAGGATTCCACGGGTGTCAGTTTACCCCCGATCGAACAAAACGTTAATCTCTGGGACTGAAACAGCCTGTCCCCCAGAGAAATTTCCAGTACCACGACCCGTCACACGACGTCGACAACAAGTCAGATTTATTTTGTCGTGATAACCTCGTTACAAACATTCTGAATTTCGGGGAATTAAACCCGTTTCCGGGCCTGGGACTTTCGTCACCGGGACCCGACGGGTTAGACGAAATGCAACACCAGGGGTCCTCGTATAGAGCGGCCGCGGCTGCTGCGGCCGTTGCTGCTTATCAAGAGCAGAACTACAGGTCAGGTGGATATCCCTTCCCCCCACAAAACCCGTACGGGTATCATCTCGGAAACTACCCGCCTCAATGTTCCTCCCCTCCGAAAGATGGTGAGTGACCGAATTTTTTAATAAGTTATTATTGTGACACGAGAATTGTAAAAATGTATTCACAGATAGTCAGGCATCGGAGAGAAATTCATTAGGGCGTACAAATTTTGGTGGTATAATGGACATTAATTGTCAGAAATTCTAAATAGTCAGATTTATTTATCGAACATTGCACTCCACAAGATCCTTTCTTATCATATTTTGTATTGCTGGTCATGATGCAATATCAATATAATCATATCACAATATATGTATAAGTCTGCATGGATTTTCTGCAGAACATTAATCCATTCTCACTCTTGTCAGTTGTAATGTTGTAAATGTATGTACACGCAAGTGTCTAAGAAAATGTGTCTCATAAAATCTCGACATTCCCCAAACAGGCATCGATGCTACCATGATTATTATTTTATCCTGGGCTTCCTTTTAAGTACTCGAATATTTTATGCATACGCTTTTTCCGGACATAATTTCTCCCATCGTGAATAATTAAGCATGTTTAACCAACTATTTAAACAAAAGCCTGGAATCATGCTCCCTACATTTATAAATATCTACTTGAGCCTAGTTAACTCTTTCACAAGGGCTGCAGGAGGGGTATCACGTATCCGACCGAATTTCCATCAGATAAACGACCACAAGAGCAAAATGGGTGCTTGCTAACTCGCTCATCCTATTAACCCGTATTACGTATTCATGATACCCGGTTGTTTGTATTCCTGGGAAAGGTTTACGAGCTCTTGTGTCACATATGTAGGAACCACCAAGGAGCTGTTGACATTAGCGGTTGGTGGGATTTCCTTCCTACGAGTGCTTAGTTAGCATGGCATGACAGTCTGTTTACTTTGGAGCTTATTACTGGGAAATATTTGAGCAAAATTGCAACTGCGAAGCCAAGAATGAAATGAAGTAACAAACCGTGAAAAGAGCAATTTTAGAGATAAAATAAAATAAAATAAATAAACGGGTTCGTATTTTTCGGCTTATGTACATATTCTTAGCTACATGTTTATGTAGTCGGCGTATTTTTGGTATTCAGCAGAACCTGAGTAAACAGAAAATAATATATAAAATAACAATGATTACGCAAAATCATTAAGTAGAAAATTTCCACGGCTCGCAGTCATAGTTGGTAAAATGAAATTGGTCTGGTTTCGTGTACAGCATGGAATAATTTAGTATGACGTGTTGAATAAGGAATTGAGGGCTCAAGTATTTCACTCTGCATTCACATTCATCATATCAAATGACACATGAAACTCTGGACAAACTGCGTACATGATCCCATCGTTAGTTGGTGAAGAACCAACCTGATCCACATCCCCCTTCCCAGCGGGACCAATAATAATAACTCGTCGCGTATGTATACCTGGTGGCCAGAATTGAATGACAATGAAACAAGGTCCTTGAGGAGTTGGGTCATGTTCGGGTCATCTGGTTTAAATGGGATCAAGTGTCAGACATACGAACTCACCCCCACCTCTCCGCCTGTACTGATCTCTGCCCTAATAAAATCGGGCATATCGTGTGCATACGTCCATTTGTGTTAAGACGGGCATAGATACGTGAAAATGGGAATAATTGTGATTTTCAGAGCCGAAAGAGGAGCCGGGTGGAATGCGGTTGAACGGAAAAGGGAAAAAGATGAGAAAACCACGGACAATTTATTCCTCCCTACAATTACAGCAGCTCAACCGCCGCTTCCAGCGCACGCAGGTTCGTTGCTTAATTATTTGGTTAATTAAACACTAATTAAAAATATGTAATAATGACTTAATTTCATTTTTGCAGTATTTAGCTCTACCTGAACGCGCGGAACTCGGTAAGAACCATCTTTATTTGCAATTAGTATTTGTAGTACAGTTTAAAATCTGGTAGAGTCACTAGTTTGGCATTAAATTCCTACCACATTGGTAGTAGTTTTGTTTCCCAGTGGTAAATTTTTCCTTATCACAACATTGCTAAACGGTAATGCCCGTGTTATTAGAGCCACATACGTACTCCGGTCAGTAATGATTTAATACCAGTTTTTTAAGAGTATATGGATAAAATATTAGCAAAAAATTCTGATCAAATTTCGATTAAGATTCCCTGTGCAATATTAATAACGCATCATTCTTATTTTATTTCAGCTGCCTCTCTTGGTCTCACACAGACACAGGTTTGCAACCCTTCCGTATTTTTGGAATTTTTACCTCATAAATACTACGAGCAATTCGGCTTGTTGTATCGCATATATACTTAAATGGTTGAGTAGATTTTACAGCAAATCATATCGTGGAGGAAAATATTATTGTATGAAATCTGAAAAGCACATCATCTCGAATATCAAAGAATTCCCTTTCGTCACCTTTTAAAAGAAGTATTAAGACTCCCAATAAATAAATCTCAAGTGCTCTGAATGTGTAAGGATCGTCATACGTGGTGCTGTTCCTCCTTCACTTCTTTTCCTCATCTTTTTCCCTCATGTCCCAATCTGCTCACCTTTCCTCACGTGAAGTTTGGCCGGAACGGAAATATCAAAGAAGCAAGACAGCCAAGATATGGGACTGAAAGAAGTTGAGAATTTAGAATCTCTTAAAGCAGCAGGGGACCATTCTTCAATTAGACACGGTTCTCCCTATTTCACTCGCCCTCTGTGCTCCACATTCCCTCATCTTCTCCTTCAGACTTATCACTTTTCGGTAGCTATTTCCTTCGTCCTAGGTGAGGTTGCAGACCTCACGAAATCTGAGACATTTCCACTTTTTACTACAAGTATGTACTCGAAAAAACAAATTTTTACACAGATTTTGAAGCCGAGAGCAATAAATTTCCTGGATAAAGATAACACAAAAACAATGCTGCTGCTCCTCTTGGTGGTAATAAAGCGAGAGATTAATTGCTTTCGGCTTGATATTTCACATTCTGTTGAGCACAGCAGTAGTTCACACAAACACAATGCGATATACGGACATAAATTCTGTTTCTAGTGAGTTATTCTCTACGTTCAAGTGTCCTTCCTGTCCCAGGTCGCACATACATAAGCATAAGTATTTAATAAATAGTAAAAAAGACAGCAGTACGGGTACCCTTTTTACTTTCCTTTTCTTAATAAAGTTCTTTTTTGCCGTAGTGTTGTACTCACCAAGTCAGAATTGAGAGCATGTCCATCCACATATCTGTAGGCAGGGAGAGGAGGAAATACAGGAACAATAGGCTGTGAGCAGGACAAGATAATTTATTTTGAGGAAGCCCGAACGGTAGCTAATCCAAGATAATTTATAAATTAGTCGTGAAGAGGGAGAAATGGAAAAAGCCTCATCATTATTTAACTTGTACAAGCAGAGAAAATCCATTGTGCTCCATTTAAAGCGTAGCGTATTAAAAGAGTCGAAGAGATGCAGCACAAGTAGTTCACTTCTTCTCCTTCTTTGGGGAAGAATGGATTAGATTTGCACGGTTGAGGTTGAGGTTGACGGTAAACAAGTGTGACATGATGTGATTGTGAGGACCTAAGTGATACCCCTCGAGCACAAGCAACCACATTTTTAACCCTTTTTTACTGTACCCCAGGAATTCCTTAAAGGATTGACAAAAAATAATGCATCGCAATTTATTTTACGATAATATTTGCGTAGGACTTGTGCACATTATGAAAATGTGGATGATGGGTGCATATTAATCGCACATGCACCCACTCCTTTCCAATGTTGTTGTTGTTTAGAATGCAGTTTGAAACAACACATTGTTGTCTGTGAGTGAGTGGAGATGGTGAATGTTTTAGCAGTTGGTAATGGAAGGGTTCTTCTACTCATTTGGTGTGGTTATTGGTGCCCCGGCCGAGTTGATGATGGGGCCGGTCTGACACCTTAGAATTAAAAGCAGCAGGGGAGCAGCTTGTCACTGTCAAAGTCTTCATGTCTTGCACTACACCAAACCAAAAGGTAATGATAAAATGCAATGTTGCGACTTTCTTCCTTCAAATGTGTGTCATGAATGAAATCCTTGGAACAAGCCATGAGACAGGGAAGTGTACATACATTCGATCGAGGGGTTCAAGTTTCTTAAACTTTTATAGAAATTGCAGAGTGGAAAGAAATGGTGAATCCCGTTGTGACGCCTTCAAAAAGTACGACTTGACTTGTTCTTGTCACATCAGAATCGAAATGAGATCAACTTTTGTAACATGGAGGGAGACGTCTGTGCTTGAATTGCTTAATCATCTCTTGGATTATTGTCCCGACGTCAGTGCGTGTGACACATGATTGTTCAAGAGGGGACTCAAGTATGGAAGTACTTGATCTCTAGGATAATCTACTCCTCCTCGTCGGAGGAACGTACTTACTTTATTATTTTGACTCCCGCCTGATGTGTGTACCGAAGGGCTTTTGTACTGCAAATTACAAATTACCTTAAATAAGTGACCTTCGTAGGCAGTAGGCACCCGCACCTTGCGGGCGTTGCATTGACCCTGTTCATTTACTAAAAGTGAAGTATTTACGAGGTTACTCCGTTCAGAGGGAAATAGGAGGACGAAAAAGTAATTGTGTTCCATTGTTCATACTGCATTCTGATCTGATCTTTTTGTATAGATTGTCAAAATATAGTATTCCAATCAGGCAGCTACTTTTGGCTTATTGCAAATTTGTTTGGGAACATTGAAATTCTAGTTTTTCCAAATTTGTGACTCGATTGAGTCGTAATTTTTTTTACAAATATAGGTAGATTTTTATTTTCACAACTGTCTCCACTTCTTGCTCCCTCGTGTACTTGGACCACTTTGAGCAAAAGTTTCCTCCTTTTCCTTTACTTTTGTGGTTTCGAGATGGTATAGGCTCGTAACTACATACGTTGCTGATGAGTGAAGTGGTGACCTTCCGAGAATAATTGAGGTATAGAAATTGAACGAGTGCTGACATTCACTTGAAAAGTGTGTATTCAAGCATTGTTCACGTTTCCTATTCTTTGATACAAGTTATACACATAGTTTCAAAACCGAAGGCCCTTCTTCGCCCATATTCACCAACCCAGCAGCATCATCTCGTCCCCGTACTGCATTGTATTCCATATTGTATTGTACAAGAGGAGAGTACATCATTTCATACAACATGAACCATAGTGAACAGCCTTTGTCTCTTTGTGCCGACTCATCTCGTCATAAACAAGAAATCTGCTGCTGCTCCTGGCTATGCATCCACGTCCAACTGACACCTTTTACCCGCAGTTTTATAAATTCTCTGAAAGGCTTCCTCAAAGTCAAGATCACTAAATATTCAGCGTTGATTTGTTACAAGTCGAGAGCGAGAGAGATGCGAGTAGTAAAGGATGGATGATGAGGTACAAAATTTCCAGTGAGTCAGTCAGTCAATTGCGGTTTCGTATTGTTGCGTAGAACTGAATGTGTCAAATCAAGATGAAGATAAAGAAAATGAGAAACAACAAGTCGACAAGGGAGCAGCAAAATACGTGGCGTGCATCATGATGCATAAGATCTTCCATTCTGGAGGAGAGTGGAAAAATTTTGGAAAATTGGACACAAAATTATGAATATTTATTTTGAAATATTTGAAATTATGATTTATTAAAAATCGTAGCAGCAACAAATCAAACACTGTTCAAAACTAAAATCTTGTGCAAATCCCATAATTGTTTTGTTGGGAATACTTTTTAAACTCGGCGAGGCTTATGAACCATTCTTCTCCTGTGTATCTGTGCATACGAAGTTTTTTAAGCCGACACCACAAATGGAACTCTTGCACAAAAGTAAAGTGATTTCATTTCGGTGCTTTGCTTTGCTTTACTTTTTTGTCGTTTTATTGACTTCGTGATGACTACAGTGGAGGGATGAATGGCGGTTGTTGGTGTTGTTGTGGACAGGAATTGAATGGTGGTGTAGTGACTCGGACACAAAACCAAGTGGAAAAAATGAATGAGTGCTAATTACAGGAAAAAGTGGAGCTTTTATTTGTCTGCTTTTTAAATGTGGTTGGTGGCATTGTGTGCTCTCTGCCACTGCCAAATTCAGCGCATCATATTTCATTGATGTTTTCCAATTTTAAATATTACTTTGAGTTCAAGTTTATAAGAAGCAATTATGAAATTACGGGATTATCAGAGATATGGTGAATTGAATTTGTGTCATTTTGATGCAAATCTCATGCCAAATAAGTTAAGACCGTGAGTAATAAACGAGATTGCTTAAATAGGCGATTGACCAGATATTTTGCAATGATTAGCACTAACACGCCGTACGAGTCATATGCAATATAAACTGAAACTTCAATTTATTTAGACAAGTATTGAGAAAAAATTCACACTTAAGCCCCACGGATTTGCATCACATGCATCCTTTTAACCCTAGGATGACCATTTTTATGTACATAAGTTATTTTAGCATTTTTTATAAAGTTGAATTTTTTTTCCCTCTGAAAGAACACAACAAGTGTTTACAAACGTGGATCTCTTATTTCTCTCGAGGCTTTTGGGAAAGGAGTGTCCCAATCCTGTCCCCGGAAGCTTCTTCGTCTTCACAACCGCCAGGGATTTCCTCGTCCTTTTCATTTCTCTCAAAATTTCCCCCAAACCCCAAAAAAATCTTATGAAACGGTAGAAGATCTACGATTAAAAATACTACGGACACCCAAACACCATAAATTTTACACTGCTATTTACTTAAATGGAACAGTATTTTTGCGTCTAGCGTGTGCATTTATTTCTGCTTAATTAGCAATCTCTGACTGGAACATTTCCACGTGTTACACAACCTCAACCCTAGGTAGACGTAGGAGTATTCGGGCGTCTCAAATAGCTCAGAGGTACATGATTTTCCAACAATACGACCTCGTATTTACAATACCGCAACCCTAATTTAATTGAACACTGTGTAAAAACACTTTTAGCTCGTACGGCACACGCTGTACAGCACCAAGCATCATTATCGTCATAAAGTTATTATATAAGTGAGTAAAAAGGAGTACGTAAACAAAGTAAACCGTGAAGATTACCAAAGAATTTTTCTCATCAAACTAATGTTTTTCAGCGAAATGAAATATATAATGGTGCACTTTTCTGTTGTACAAAGTCAAATTTTATGCTACACTTGCTTGCATTTACTCATTTTAGGTAGTGGTAAGCAAGGGAGGAATGTGCAGCATGCTGTAGTAAAATCGCATCCATTCATTCCTCCATTCCACTACTTGTTGGAGTACACTTAATTGTTTAATTGTTGAAGGTATTGTAGGAGATTTAGAGATACATATTCAAGATGCGAGATGAGTTATTTCAGACGTAAAGGAATCTGACAACGATATGTATGTGCATATGTGCATATGAAAAGTGTGGAAGAACCTGCACGATGACTTCTCATTTATTACATGCTCCGTGTAACGAGTAATTTCTATTTTCCGTACTATTATTATTATGGATGTAAGCTAAACTATCACTAATTTATTGCATAGGTACAACGCGATAGTGTTGTTCCTAATGCAAAATTACAATGTAGAATGGTCATAAATTTAGAGTCAAAATTCACTTCATGCTCGTTCCTCATGCTCGTACGTACCTACGTAGGTTCCTTATAAATTCGCGTGTGTGTGAATGGGTTTTCTTTTCACAGAGGCACATGAGTTATCGATTTCTGCAAAATTTTTTAGTGGATCAGATAACGCATAAATGGTGTTATCGTCACACTACGCCAAAACTGCTACCAGCTGGTGTACGTGTGCACATACGAGTTTATTCCATCCTGCATAATTCTACTCGCTGACCAATTGACCATTATTAAGTACTATTTTGCATTTTGGGTTGTTTCTTGGCAAGCTATTTGAGGTAAAAGTCCTTGTCAACGTGGTCGTCTTGTGGTGGTTGGTGTGCAGTGGTGGGTTCCATGCCTTGGTGCGTGTGGTTGTGGGCGAAGGCTGTCGTTTACGCTATCCGTGACATGTGGCCAGATTGGTTATGCGGTTTTCGCAATTTGTGGAATCGTGGCAAAAGAACAACAACAATGCGAAACGAGATGCAGTGCAATATCGATCACTTAAAATAAAAAAGTTGTGTGGGTGACATTGTTTTTTATTATCTTGTATTTTTTCTTCTTTCATGCGCACGCACAAGGTTAATATGTAAAATCATAGATATGAAGGGAATTTAGACGAAAACTGCATGCATGCACAAACGTGATGGTTGGTAAATATCTTAAAAAAGTTGTGCTCAGTTTATGATGATCAGACACAAGCTGAAATAATACGAAATGAAGGGAAATGAAACTTTTTAGAGCTAAGACTACATCCCTTCCTTCGCTCGCTTCGAATACGGCCAGTCTGTCAGTAACCAACTGTTTTATCTGGTCTGGCCTTCTTGTCAATCATCATCTCTTACTTTGCTTACCACATGCATTACCTGTTCTCTCGTGTTTTGTGCTATACTAAAATGCAGACTTTTCTATATGATATTCTCACTAAAGCTGCAACATACTTCGTTCCTCGCTGGCTTACATTCACATCACATCACAACATGGCATAATATGATGATGGCAGTAACAAAGGCTGCCAAGCTGCTTCATTTCATTTTATTACTTGTACAATTCCACACCACGTAAATGCATGCATGTAAGTAAGCACCTACTTTTACTTATTTACCACTCGTAAAATAATGCCGATTGGGTGCAGGTACCCACATTTTCGAGTGGCCCTAACAGCCCCATGGATCGAGTAAAGAACACAATTTTGGACATTATTGATTTTGCATGTACACAAATAAATAGGTTCGTTCGTGTTGCAAAATTGAGCTATTTATAATGTAATGTATCTTGTGTCGGATAATTAAAAATGAGATTACCTTTAAAGTAAACATGTAAATAGGGTTGATGAAAAAAGTCATATCCCTCACGGTCCTTGTGTCAGGTTGGTCGGCCCACATTAATTCAATCCCACATTATCCTCGCATCATAGAAAATGAGAGATATCTATGAGTGGAAAAGAATGGAAATATGTGAGTACTACGCGTTTTGTTTGATATGCATGTAGAATAGGTGCGATGTGTGATGGATGGATGGAAGAAAGAGGAGTGTGTAGAGATCATATAAAAGTTTTTAAGCAGATACAAGTACGTATTTGATGCATGTTATGCTCGGACCGGCCCAAACAGCAACCAGCTTGAGATTTTGGTTTGAAATTTTGAGATCAATCTGACAGCTTTTGATGTGGGTTGTCCCAACTCGAGTTAGTTTTCAGGCCATGGGAGATGCATCTCGTCATGATGATCATCAAATCATACCACCTCGCTCGTAAATACAAAGGCCAAACACGTACTCCTTGGATGGATGGAATCCTTCTCGAAGGTTGCTCCATCCCTGCAACACAAAATTTAAATGAATTTCAAGCATTTAGATTAAAATAAAATAATTAATTTCATCAAGAGTATCGTGAAGTTGCGCAATACGTAACTCACTCAGCTCCATTCTACTAAGGAGTATGTACAACTTGATTAAGAATTGCTTACATGATTCGATTAAATTGGAGAAAATGGAGTTGAGCAAATTTAATGTATCAACTGCTCATTATTGTGTAAAATAGTTGACCCAAGAACAGCATTAGATCAGCCATGAATAGGGTAGTACCACCACCAGACAACCAAATATAACAGCTCATTAGACGTCAATGTAGCAGAAAATCTCAGCCGGCTTATTTACCAAGTTACAAATGAATCTCACAAACAAATCCGTAAATCTTGTTGAGCTCTTTCTTCCTTTACCGGTACTTATCCTTAGCCAGAGTAAAGGGACCTGAGGGCCAGACTGGGCGAAAAAGCTTCTTCTTCATCTCGTAACAAGGGCCGACAGCAGCAGGCCAAAAGTTATACAACCTTTATTTTTATTACCAAACCAAACCAAATGAACCGACGGTCGACCACCGCTGCATACCTACCAGCACCTTTATCTACAATGCGATAATACTGCATGTAACAACTAACAAACCGCTAATTGATTTAAATGAATCTTATTCTTAAAGACTTGCTGAACTCGTCGATCTCATCATTCCATTGTTTGCGCAAAAGTTACATAAATACAATTGTAAGATGGAAAAATGATTAAAACAACAAAGTCAAAAGTTGTAAAGATTATCGGAGCACGTATCAATTCATAACCTGACTAATCAACATTTTCACCTTAAATGAGCTATACAGGAAGCTGAGGAGTTGGTTTAGCTGGGGGCACTTTTTGGTCATATGTTCATACTTTTGTGCAAATGACTTTATTCTCCAGTCCTGAAAAAGAAGAGCGTATTCGTTCGTTAGTTAGTTGGTCATCAGGAAGCCTTGCATTATTTTATTACGACCCGCATCATCTCTTTTTTAATAAATTAGTCGTACTTTGTTAGCTGCGTGGCGTGTGTGTGTGGATTATGGTAGTGGCAATAAATTATGGAAAAGCTTAGAAAAATTATTTCCAAAGAACTTCCTCTCTTTATTCCGAAGACTTCCTCAGGCAAATTTTTTCTCGCAGTTCGAAATGCGGAGGAAAACATTATTTTTAGATTAGCATCTCAACTCGCTGTCCCATACGCTTACCTGCCTTCGTATGTGTACCCTTTCCTTGAGAGTAAGGCCGAGTGGGCGTGGGAGGCGTTGTGCTGCAAGTTGGTCGATGTATTTAGATGGCAAAGTTCTCCGTAGGGAAGGTAAGGGTTGCTTTTTACACGGCAGAAGAGTCTCCTCGCATTGTCGGAAGAATAACACTTTGTAGCAATAATATACATAATGTGGACGTAATGCACGGTCAACGGTCCGCCGAACGAGAGTCTCCTCTTAATGATGATCATGTGAACGTACGTAAACATGTACCACGGGTCGGATTTTTTGTGAATTTAAGGTAAAAGTAAAAATATCGTAATCAGGGAAGAGAAATCTGAACGCATGGAATAAAAAATTAGGGTTAAGGGTTAAGATTTCGTGTGGTGGTGTGCGTCTGCCTACATGATCCTGCCTAATGTGGTGGTATGGC

Red:

PFE

Gray:

sequence confirmed by PCR

5’ Genomic region

Putative

Exon1

Putative

Exon2

Putative

Exon3

Putative

Exon4

>FCD Dll full length

ATGCAACACCAGGGGTCCTCGTATAGAGCGGCCGCGGCTGCTGCGGCCGTTGCTGCTTATCAAGAGCAGAACTACAGGTCAGGTGGATATCCCTTCCCCCCACAAAACCCGTACGGGTATCATCTCGGAAACTACCCGCCTCAATGTTCCTCCCCTCCGAAAGATGAGCCGAAAGAGGAGCCGGGTGGAATGCGGTTGAACGGAAAAGGGAAAAAGATGAGAAAACCACGGACAATTTATTCCTCCCTACAATTACAGCAGCTCAACCGCCGCTTCCAGCGCACGCAGTATTTAGCTCTACCTGAACGCGCGGAACTCGCTGCCTCTCTTGGTCTCACACAGACACAGGTGAAAATTTGGTTTCAAAATAGGCGGAGCAAGTACAAGAAGTTGATGAAGGCAGCTCAAGTGCCCGGAGGGTCGAATCAACCGTCGAATCAGCAGGGAGAAAATTCGAATGAGACGATGAGTCCTCAGCCCCCAGATTCTTTCCCTTCGCAACCCGGTGACCTCAGCCCGCCTCCAAATCCTAACCCCCAGTCTCAGGGAGGTTGTTCATCGCCCGTGTCGCCGTGGGATATAAAGGGTGGCGGCGGGGGCGGGGGTCCTCAGCATCAAAACGGTCCCCCCAACATTCCTCAACTTCCCCACCACCCACACCCCGGAATCCCCGTAGGGCACCAACCTTATCAATATCACTGGTACCATCAGGACCACTCCCTTCTCACGTAACTAACACACCTCAACTCTAA
